# Brain-Engrafted Monocyte-derived Macrophages from Blood and Skull-Bone Marrow Exhibit Distinct Identities from Microglia

**DOI:** 10.1101/2024.08.08.606900

**Authors:** Siling Du, Antoine Drieu, Yumeng Cheng, Steffen E. Storck, Justin Rustenhoven, Tornike Mamuladze, Bishan Bhattarai, Simone Brioschi, Khai Nguyen, Feiya Ou, Jay Cao, Patrick F. Rodrigues, Igor Smirnov, David DeNardo, Florent Ginhoux, Marina Cella, Marco Colonna, Jonathan Kipnis

**Affiliations:** Department of Pathology and Immunology, Washington University in St. Louis, School of Medicine, St. Louis, MO, USA; Brain Immunology and Glia (BIG) Center, Washington University in St. Louis, School of Medicine, St. Louis, MO, USA; Université Paris Cité, Institute of Psychiatry and Neuroscience of Paris (IPNP), INSERM U1266, Paris, France; Center for Brain Research, Faculty of Medical and Health Sciences, University of Auckland, Auckland, New Zealand; Institut National de la Santé Et de la Recherche Médicale (INSERM) U1015, Equipe Labellisée—Ligue Nationale contre le Cancer, Villejuif, France

**Keywords:** Microglia, Monocyte, Monocyte-derived Macrophages, Skull Bone Marrow, Macrophage ontogeny

## Abstract

Microglia are thought to originate exclusively from primitive macrophage progenitors in the yolk sac (YS) and to persist throughout life without much contribution from definitive hematopoiesis. Here, using lineage tracing, pharmacological manipulation, and RNA-sequencing, we elucidated the presence and characteristics of monocyte-derived macrophages (MDMs) in the brain parenchyma at baseline and during microglia repopulation, and defined the core transcriptional signatures of brain-engrafted MDMs. Lineage tracing mouse models revealed that MDMs transiently express CD206 during brain engraftment as CD206^+^ microglia precursors in the YS. We found that brain-engrafted MDMs exhibit transcriptional and epigenetic characteristics akin to meningeal macrophages, likely due to environmental imprinting within the meningeal space. Utilizing parabiosis and skull transplantation, we demonstrated that monocytes from both peripheral blood and skull bone marrow can repopulate microglia-depleted brains. Our results reveal the heterogeneous origins and cellular dynamics of brain parenchymal macrophages at baseline and in models of microglia depletion.

## INTRODUCTION

Microglia, the major immune cell type situated in the central nervous system (CNS) parenchyma, account for 5-20% of total glial cells in the CNS^1,2^. During CNS development these cells play important roles in various processes such as neurogenesis, synaptic elimination, and the establishment and rewiring of neural circuits^3–7^. Fate-mapping studies using *Runx1^MER-Cre-MER^* and *Csf1r^MER-Cre-MER^* mice have demonstrated that microglia originate from yolk sac (YS) progenitors, and their development is independent of MYB, the transcription factor essential for the development of hematopoietic stem cells (HSC)^8,9^. Under normal conditions, studies using pulse-chase labeling, parabiosis, and bone marrow transplantation have shown that microglia predominantly maintain themselves through self-renewal, with minimal monocyte-derived macrophages (MDMs) infiltration from the bloodstream, reinforcing self-renewal as the primary mechanism for microglia maintenance throughout life^10–14^.

The profound self-renewal capacity of microglia, however, presents challenges in developing cell therapies targeting neurological disorders. Dysfunctional microglia are implicated in various neurological conditions, spurring efforts to develop efficient microglia replacement strategies^15–18^. Traditional approaches often involve irradiation, known to compromise the blood-brain barrier and trigger cytokine storms^19,20^. Our research, alongside studies by others^21,22^, has shown that genetic models of microglial loss can induce MDM engraftment independently of irradiation, paving the way for less-invasive methods of MDM engraftment through microglia depletion. However, pharmacological depletion of microglia in wild type mice showed no evidence of MDM infiltration^12^. Furthermore, recent studies using a repetitive microglia depletion model showed prolonged microglial loss, but the possibility of MDM infiltration in such contexts has not been thoroughly examined^23,24^. Our study aims to investigate whether prolonged microglia depletion could serve as a viable model for microglia replacement by MDMs independent of irradiation.

In pathological conditions such as traumatic brain injury, stroke, and neuroinflammation, substantial brain engraftment of MDMs has been observed, with brain-engrafted MDMs acquiring distinct characteristics from YS-derived microglia^25–29^. Notably, clonal hematopoiesis of brain macrophages has been associated with protection from Alzheimer’s disease (AD) in humans, suggesting a role for peripheral myeloid cells in neurodegeneration^30^. Understanding the mechanisms underlying peripheral cell infiltration and the distinct biology of peripherally-derived cells versus YS-derived microglia is crucial for elucidating the heterogeneity of brain myeloid cells and their roles in health and disease.

Additionally, many studies employ parabiosis mouse models to demonstrate the absence of brain-engrafted MDMs^10,12,14,31^. Although parabiosis effectively achieves chimerism in the blood for tracing blood-derived cells, it results in significantly lower chimerism within the skull bone marrow. Our recent studies have revealed that the skull bone marrow can supply various immune cells, such as monocytes, neutrophils and B cells, to the cranial dura^32,33^. However, it remains unclear whether skull bone marrow could serve as a potential source for brain parenchymal macrophages.

To address these questions, we utilized lineage tracing tools for MDMs to investigate their presence in the brain at baseline and in models of microglia depletion. Our studies demonstrated that MDMs are capable of engrafting into the brain following repetitive microglia depletion, exhibiting a distinct cellular signature compared to YS-derived microglia. Furthermore, using parabiosis and skull transplantation models, we discovered that MDMs from both the blood and skull bone marrow can replenish the brain macrophage pool in scenarios of microglia loss.

## RESULTS

### Monocyte-derived macrophages (MDMs) are present in the homeostatic brain and increase after microglia depletion

Given the selective and transient expression of *Ms4a3* in granulocyte myeloid progenitors^34^, their progeny monocytes and neutrophils are efficiently labeled with tdTomato in the *Ms4a3^Cre^:R26-tdTomato* mice (Fig. S1A). Therefore, this model can discriminate between monocyte- and embryonic-derived macrophages based on their tdTomato expression. To investigate the presence of brain-engrafted MDMs during microglia turnover, we fed mice with the pharmacological inhibitor of colony-stimulating factor-1 (CSF1R), PLX5622, to induce microglia loss with either one dose of the drug (PLX-1x)^35^ or repetitive treatment (PLX-3x; Fig. 1A)^23,24^. The repetitive PLX-3x treatment has been demonstrated to result in prolonged microglia depletion that requires a longer time for reconstitution, although the source of the repopulated microglia was not identified ^23,24^. Understanding the sources of microglia both at baseline and in the context of microglia depletion is crucial for elucidating the mechanisms behind microglia regeneration and their functions. Flow cytometry revealed that a few tdTomato^+^ MDMs (0.19% ± 0.15) can indeed be found in the microglia pool at baseline (Fig. 1B-C; gating strategy shown in Fig. S1B). Whereas PLX-1x treatment resulted in only a slight increase in tdTomato^+^ MDMs (1.36% ± 0.26), repetitive treatment (PLX-3x) led to a dramatic rise in tdTomato^+^ MDMs (47.49% ± 4.54) found in the brain one month after repopulation (Fig. 1D, Fig. S1C). The notable increase in tdTomato^+^ MDMs observed after PLX-3x treatment could be attributed to the engraftment of peripheral monocytes or the expansion of resident MDMs present at baseline. To determine whether tdTomato^+^ macrophages are resistant to PLX treatment and therefore expand during the repetitive treatment, we harvested the brains of *Ms4a3^Cre^:R26-tdTomato* mice at the end of the PLX treatment (Fig. S1D-F). Flow cytometry analysis revealed that none of the few remaining microglia were tdTomato^+^, confirming that tdTomato^+^ brain parenchymal macrophages present at baseline were also depleted after PLX treatment. Thus, the observed increase in tdTomato^+^ brain parenchymal macrophages post-PLX-3x treatment is likely due to the peripheral cell infiltration. To further track the timeline of MDM brain engraftment, we also harvested the brain one week after depletion (Fig. S1D-F). Although the brain macrophages had not fully repopulated yet, we already observed tdTomato^+^ MDMs at this time point, suggesting that the infiltration of MDMs into the brain occurs within one week of PLX removal. This early presence of tdTomato^+^ MDMs underscores the rapid response and migration of peripheral monocytes into the brain following microglial depletion.

**Figure 1.**
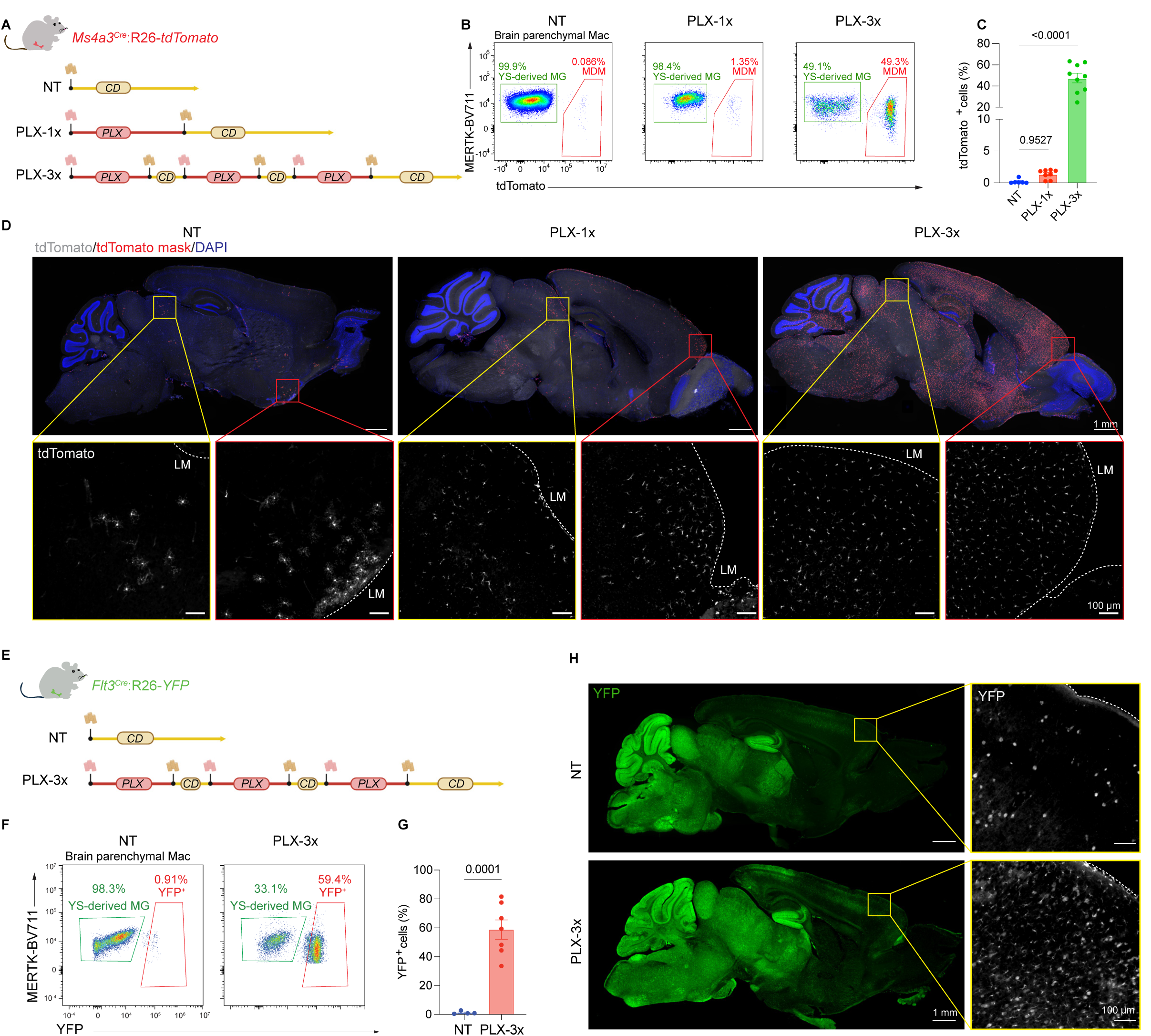
Monocyte-derived macrophages (MDMs) are present in the homeostatic brain and increase post-microglia depletion and repopulation. (A) Scheme of PLX treatment on *Ms4a3^Cre^:R26-tdTomato* mice. The *Ms4a3^Cre^:R26-tdTomato* were fed either a control diet (CD; non-treated NT), a single PLX treatment for 2 weeks (PLX-1x), or repeated PLX treatments (3 cycles of 2 weeks on followed by 1 week off; PLX-3x). (B) Representative flow plots of brain macrophages from control (NT) or PLX-treated *Ms4a3^Cre^:R26-tdTomato* mice. The percentage of tdTomato^+^ cells over all brain parenchymal macrophages are shown. The brain parenchymal macrophages were determined by pre-gating on: Singlet, Live, CD19^-^, CD45^+^ CD11b^+^, Ly6G^-^, Ly6C^-^, Mertk^+^, CD11c^-^, CD206^-^ cells. The detailed gating strategy is shown in sup. Fig. 1B. (C) Quantification of the flow data showing the percentage of tdTomato^+^ macrophages over all brain macrophages. (NT n=6, PLX-1x n=8, PLX-3x n=9; 2-6 months old). Analyzed by one-way ANOVA test. Data pooled from three independent experiments. (D) Upper panels: immunofluorescent images showing tdTomato^+^ cells in the brain of NT, PLX-1x, PLX-3x treated *Ms4a3^Cre^:R26-tdTomato* mice (tdTomato^+^ cells are masked in red, and the tdTomato signal is shown in grey), scale = 1 cm; Lower panels: enlarged images of the selected regions, scale=100 μm, dashed white lines indicate the leptomeninges (LM). (E) Scheme of PLX treatment on *Flt3^Cre^:R26-YFP* mice. The *Flt3^Cre^:R26-YFP* mice were fed either a control diet or repeated PLX-3x. (F) Representative flow plots of brain macrophages from control (NT) or PLX-treated *Flt3^Cre^:R26-YFP* mice. The percentage of YFP^+^ cells over all brain parenchymal macrophages are shown. The YFP^+^ gate was determined based on a WT mouse. (G) Quantification of the flow data showing the percentage of YFP^+^ macrophages over all brain macrophages. (NT N=4 mice, PLX-3x N=7 mice; 4-6 months old). Analyzed by parametric unpaired t-test. Data pooled from two independent experiments. (H) Left panels: immunofluorescent images showing YFP^+^ cells in the brain of NT and PLX-3x treated *Flt3^Cre^:R26-YFP* mice, scale = 1 cm; Right panels: enlarged images of the selected regions, scale=100 μm, dashed white lines indicate the leptomeninges (LM). The YFP^+^ cells present in the NT brain are neurons.

The distribution of the brain-engrafted tdTomato^+^ MDMs varied across different treatment conditions (Fig. 1D). Control *Ms4a3^Cre^:R26-tdTomato* mice exhibited scarce tdTomato^+^ MDMs, which were found at the superior colliculus and the ventral striatum. Interestingly, tdTomato^+^ MDMs resided near the leptomeninges and exhibited the highly ramified morphology associated with homeostatic microglia (Fig. 1D). The tdTomato^+^ cells in the brain also expressed IBA1, validating their macrophage identity (Fig. S1C). Following PLX-1x treatment, an increase in tdTomato^+^ MDMs was noted in the superior colliculus and frontal cortex. Post-PLX-3x treatment, a drastic increase in tdTomato^+^ MDMs was observed throughout most brain regions, though they were less frequently found in the olfactory bulb (Fig. 1D, Fig. S2A-C). PLX-3x treatment also led to a notable presence of MHCII^+^ IBA1^+^ brain parenchymal macrophages, particularly abundant in the olfactory bulb (Fig. S2A-C), indicating heterogeneity in the sources and phenotypes of repopulated microglia across different brain regions. To further validate our findings, we employed the *Flt3^Cre^:R26-YFP* lineage tracing model, which labels cells derived from hematopoietic stem cells^36,37^ (Fig. 1E). In control animal, only a few brain macrophages were YFP^+^ (1.22% ± 0.52). However, following PLX-3x treatment, YFP^+^ MDMs constituted approximately 50% of the brain macrophages (58.74% ± 6.71) (Fig. 1F-H). This significant increase post-treatment confirmed a substantial recruitment of peripherally derived MDMs into the brain. In summary, our data demonstrate that while MDMs are present in very low numbers at baseline, their engraftment into the brain is markedly enhanced after prolonged microglial depletion, accounting for about half of all brain parenchymal macrophages.

### Brain-engrafted MDMs obtain distinct cellular and transcriptional signatures from YS-derived Microglia

Using the PLX-3x paradigm, we have developed a model that facilitates MDM brain engraftment and allows us to simultaneously study YS-derived microglia and MDMs within the same environment. To further explore the cellular features of brain macrophages from different sources, we performed Sholl analysis on PLX-3x-treated *Ms4a3^Cre^:R26-tdTomato* mice (Fig. 2A-B). Notably, the branches of tdTomato^+^ MDMs were shorter and less complex than those of tdTomato^-^ IBA1^+^ YS-derived microglia (Fig. 2C), indicating a more ameboid and activated microglial phenotype^38,39^, which is reinforced by their trending increased sphericity (Fig. 2D). Additionally, we examined the expression of CD68, a lysosomal glycoprotein upregulated in activated phagocytic microglia^28^. Compared to YS-derived microglia, brain-engrafted MDMs showed higher CD68 volumes, suggesting that MDMs are more activated (Fig. 2E).

**Figure 2.**
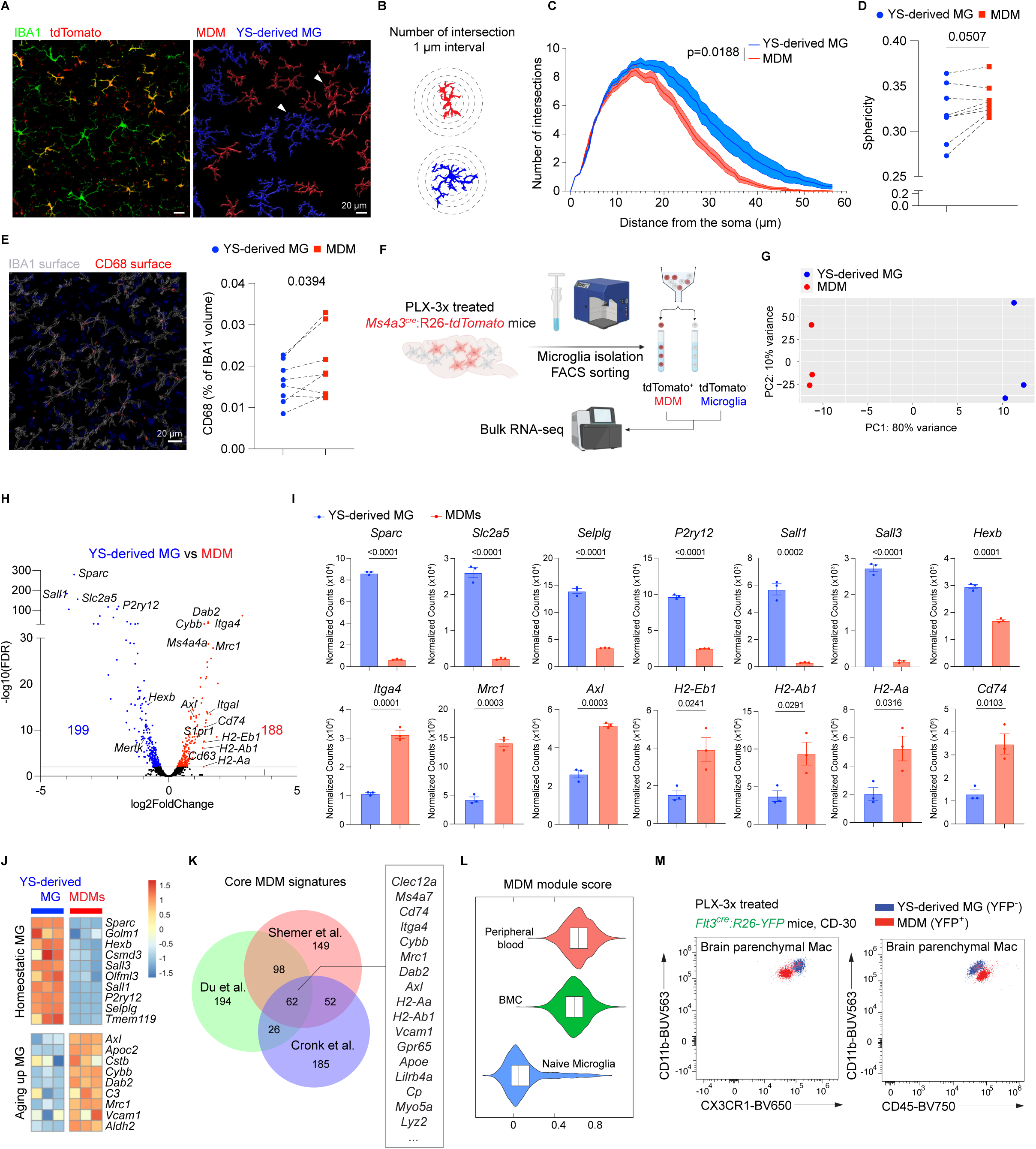
Brain engrafted MDMs obtain distinct cellular and transcriptional signatures from YS-derived Microglia. (A) Left panel: representative confocal images of *Ms4a3^Cre^:R26-tdTomato* mouse after PLX-3x treatment, IHC of tdTomato^+^ (red) and IBA1^+^ cells (green); Right panel: MDM (IBA1^+^ tdTomato^+^) and YS-derived microglia (IBA1^+^ tdTomato^-^) surface masks were determined by Imaris. (B) Schematic illustration of Sholl analysis. (C) Sholl analysis of YS-derived microglia (YS-derived MG) and MDMs (n=9 mice), 15-25 cells per cell type for each mouse. Analyzed with two-way ANOVA with Geisser-Greenhouse correction and Sidak’s multiple comparisons test. (D) Quantification of the sphericity of YS-derived MG and MDMs determined by Imaris (n=8 mice). Parametric paired t-test. (E) Left panel: representative confocal image with Imaris-masked surfaces of IBA1 (white outline) and CD68 (red) in PLX-3x treated *Ms4a3^Cre^:R26-tdTomato* mice. Blue: DAPI. Scale = 20μm. Right panel: quantification of the CD68 proportion per macrophage. The macrophage volume is determined by the IBA1 surface (n=8 mice). Parametric paired t-test. (F) *Ms4a3^Cre^:R26-tdTomato* mice were treated with PLX-3x and microglia were isolated for FACS sorting followed by bulk RNA-sequencing. (G) Principal component analysis (PCA) of transcriptomes of MDMs (tdTomato^+^) and YS-derived microglia (tdTomato^-^). (H) Volcano plot showing differentially expressed genes (DEGs) for each group. Genes labeled in blue (enriched in YS-derived MG) or red (enriched in MDMs) have a log2FC>0.5 and FDR<0.01. The number of upregulated and downregulated DEGs was indicated in corresponding color. Sequencing data was analyzed by DESeq2, and a complete list of DEGs is included in Table S1. (I) Normalized RNA counts for select marker genes, analyzed with unpaired t-test. (J) Heat map of representative genes associated with homeostatic microglia^70^ and aging microglia^71^. Each row is z-score-normalized counts for each gene. (K) Venn diagram of DEGs in MDMs signatures identified in the current study, Shemer et al.^45^, and Cronk et al.^21^. The data were all analyzed via DESeq2. A complete list of the core MDM signature genes, and the gene lists used to generate the plot were included in Table S1. (L) Violin plot of the geneset score of core MDM genes on different cell population isolated from Xu et al.^46^. The AddModuleScore() function from Seurat was used to determine the geneset score. (M) Representative flow cytometry plots of CX3CR1 and CD11b expression (left panel), or CD45 and CD11b expression (right panel) of YFP^-^ (blue) or YFP^+^ (red) brain parenchymal macrophages of the *Flt3^Cre^:R26-YFP* mice with PLX-3x treatment. CD-30 represents the mice treated with common diet (CD) for 30 days after PLX removal.

The morphological disparities between brain parenchymal macrophages derived from different sources suggest that they may possess distinct transcriptional signatures. To assess the transcriptional differences between brain-engrafted MDMs and YS-derived microglia, we isolated tdTomato^+^ MDMs and tdTomato^-^ YS-derived microglia from *Ms4a3^Cre^:R26-tdTomato* mice one month after PLX-3x treatment for bulk RNA-sequencing (Fig. 2F). Principal component analysis showed that tdTomato^+^ MDMs are dramatically distinct from YS-derived microglia (Fig. 2G). Differentially expressed genes (DEGs) enriched in YS-derived microglia included homeostatic microglia genes such as *Sparc*, *Slc2a5*, *P2ry12*, *Selplg*, *Hexb* and transcription factors *Sall1*, *Sall3* (Fig. 2H-I, Fig. S2D, Table S1), whereas brain-engrafted MDMs showed increased expression of integrin *Itga4*, mannose receptor *Mrc1*, and phagocytic receptor *Axl*. In addition, brain-engrafted MDMs expressed higher levels of MHCII-associated genes such as *H2-Eb1*, *H2-Ab1*, *H2-Aa*, and *Cd74* (Fig. 2H-I, Fig. S2D, Table S1). These DEGs indicated that MDMs may be more actively engaged in cell adhesion, pathogen recognition, and phagocytosis. To further understand the transcriptional differences between YS-derived microglia and MDMs, we performed gene set enrichment analysis (GSEA) to determine if the observed gene expression differences between YS-derived microglia and MDMs overlapped with previously published gene sets of microglia under various conditions^29,40–43^ (Fig. S2E). Notably, we observed a higher enrichment score for bone marrow (BM) cells engrafted to the *Csf1r*^-/-^ brain via intracerebral transplantation (ICT) in the MDMs compared to resident microglia^29^. MDMs also displayed significant enrichment in gene sets associated with microglial immaturity, response to LPS challenge, aging, and AD from prior studies^40–43^. In contrast, YS-derived microglia showed higher enrichment for homeostatic microglia, microglia transplanted to the *Csf1r^-/-^*brain, and microglia maturity (Fig. 2J, Fig. S2E). In addition, GSEA using the molecular signature database^44^ revealed an enriched signature of the complement pathway and inflammatory response in MDMs, indicative of a more activated state. In contrast, YS-derived microglia displayed higher enrichment in apical junction pathways, suggesting a more resting and homeostatic state (Fig. S2F).

Next, we aimed to create a general gene list to assist the identification of brain-engrafted MDMs in various transcriptomic studies. To achieve this, we reanalyzed two public bulk RNA-seq datasets that profiled brain-engrafted bone marrow cells (BMC) after irradiation and BM transplantation^21,45^. We then overlapped the signature genes of peripherally derived macrophages from these datasets with the MDM signature genes identified in our study to generate a core signature for brain-engrafted MDMs. We identified 62 genes shared across all three datasets, including *Clec12a*, *Ms4a7*, *Cd74*, *Itga4*, *Cybb*, *Mrc1*, *Dab2*, and *Axl* (Fig. 2K, Table S1). To test the robustness of our core MDM signature in a different transcriptomic study, we conducted a module score analysis using our core MDM signature on a published scRNA-seq dataset containing naïve microglia, microglia replaced with BM cells, and microglia replaced with peripheral blood cells^46^. Indeed, microglia that were replaced with BM cells and peripheral blood exhibited significantly higher MDM scores compared to naïve microglia (Fig. 2L).

Lastly, to validate the transcriptional differences between YS-derived microglia and MDMs at the protein level, we conducted flow cytometry on PLX-3x treated *Flt3^Cre^:R26-YFP* mice. We observed that brain-engrafted MDMs possessed lower expression of the fractalkine receptor CX3CR1 and integrin CD11b, which are involved in their surveillance in homeostasis (Fig. 2M). Unlike YS-derived microglia, which express an intermediate level of CD45, brain-engrafted MDMs exhibited higher CD45 expression (Fig. 2M). In line with the bulk RNA-seq data, we observed lower expression of Mer tyrosine kinase (MERTK), and higher expression of F4/80 in MDMs by flow cytometry (Fig. 2M, Fig. S2G-H). Thus, our results show that YS-derived microglia and MDMs not only differ morphologically, but also exhibit substantial transcriptional and protein-level variations which could potentially influence their functionality in the brain environment.

### Brain-engrafted MDMs transiently express CD206

From our bulk RNA-seq analysis, we discovered that MDMs upregulated *Mrc1* compared to YS-derived microglia (Fig. 2I). Mrc1 encodes for mannose receptor CD206, which is highly expressed by dura macrophages and parenchymal border-associated macrophages (PBMs) in the leptomeninges and perivascular space, but not typically expressed by adult microglia^13,47–49^. Intriguingly, CD206 has also been found on precursors of CNS-associated macrophages. CD206^+^ A2 cells derived from the YS can infiltrate both the meningeal space and the brain parenchyma, subsequently losing their CD206 expression and acquiring a microglia phenotype after entering the brain^49,50^, while the meningeal cells retain their CD206 expression and further differentiate into various subsets of meningeal macrophages^13,51^.

To assess whether MDMs undergo a similar transient expression of CD206 upon brain engraftment as during development, we generated *Mrc1^CreERT^*^2^*:R26-tdTomato* mice to enable transient labeling of Mrc1-expressing cells (Fig. 3A, Fig. S3A). After tamoxifen injection in control animals, over 50% of dura macrophages and PBMs were labeled, while no recombination was found in microglia and blood monocytes (Fig. S3B-D). By immunohistochemistry (IHC), we observed tdTomato^+^ cells exclusively in the perivascular space and leptomeninges, co-localized with the CD206 staining (Fig. 3D). However, in mice treated with PLX-3x to promote MDM engraftment, tdTomato^+^ cells were also found in the brain parenchyma, co-localizing with IBA1 (Fig. 3D, Fig. S3E). These tdTomato^+^ brain parenchyma macrophages also expressed high levels of CD45, while exhibiting low expression of CX3CR1 and CD11b, indicative of their MDM identity (Fig. S3F). Notably, the tdTomato^+^ cells were CD206^-^, suggesting that CD206 expression on the cells is transient (Fig. 3D). To test whether CD206 expression in brain parenchymal macrophages is downregulated over time, we harvested brains at 7, 15, and 30 days after PLX-3x treatment (CD-7, CD-15, CD-30) and assessed CD206 expression by IHC. Indeed, we found a substantial decrease in CD206 expression, suggesting that after engrafting the brain, CD206 expression gradually decreases in macrophages (Fig. 3E, Fig. S3G-H).

**Figure 3.**
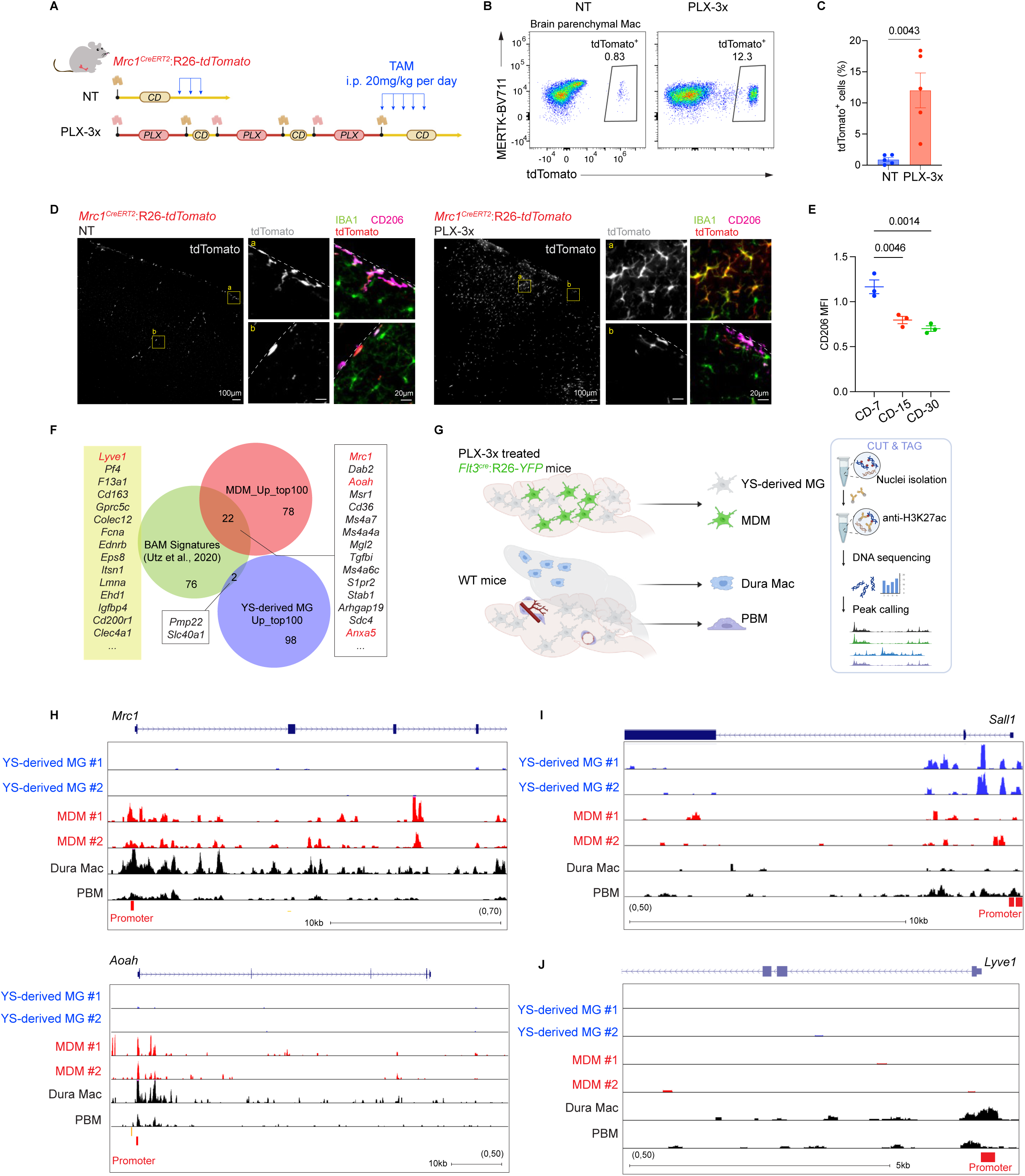
Brain engrafted MDMs transiently express CD206. (A) Scheme of PLX treatment on *Mrc1^CreERT^*^2^*:R26-tdTomato* mice. The *Mrc1^CreERT^*^2^*:R26-tdTomato* mice were fed either a control diet (NT) or PLX-3x. Upon the removal of PLX, the mice were injected with 20mg/kg tamoxifen for 5 times at D0, D1, D2, D3, D5 and D7 to enable the transient labeling of brain engrafted MDMs. (B) Representative flow plot of tdTomato^+^ brain parenchymal macrophages from different treatment conditions. (C) Quantification of the flow data showing the percentage of tdTomato^+^ macrophages over all brain macrophages. (NT n=5, PLX-3x n=5). Analyzed by parametric unpaired t-test. Data pooled from two independent experiments. (D) Immunofluorescent images showing tdTomato^+^ cells in the brain of control (NT; left panels) and PLX-3x treated (right panels) *Mrc1^CreERT^*^2^*:R26-tdTomato* mice. For the larger images, tdTomato signal is shown in grey, scale = 100 μm. The enlarged images of the selected regions showed the single channel images of tdTomato (in grey); and multiple-channel images of IBA1 (in green), CD206 (in magenta) and tdTomato (in red), scale=20 μm, dashed white lines indicate the leptomeninges. (E) Normalized mean fluorescent intensity (MFI) of CD206 analyzed from confocal images of PLX-3x treated mice of different time points after PLX removal. CD-7, CD-15, CD-30 represent the mice treated with common diet (CD) for 7, 15 or 30 days after PLX removal respectively. N=3 mice for each time point, data acquired from a single experiment. The CD206 MFI was normalized to the MFI of DAPI for individual image. (F) Venn diagram of the top 100 genes from the BAM signatures identified in Utz et al.^47^, and the MDMs signatures and the YS-derived microglia signatures identified in the current study. The data were both analyzed via DESeq2. The gene lists used to generate the plot were included in Table S1. (G) Scheme of H3K27ac CUT&TAG on different macrophage populations. *Flt3^Cre^:R26-YFP* mice were fed PLX-3x, then changed to common diet to isolated YFP^-^ YS-derived microglia and YFP^+^ MDMs from the brain. The dura macrophages and PBMs were isolated from WT mice by FACS. Cells were then processed with H3K27ac CUT&TAG. (H-J) CUT&RUN analysis of H3K27ac peaks at promoter regions of genes *Mrc1*, *Aoah*, *Sall1* and *Lyve1*. The snapshots of UCSC genome browser are shown, and the promoter regions are highlighted in red.

Given that the *Mrc1^creERT^*^2^*:R26-tdTomato* mice also labels meningeal macrophages, it is possible that the tdTomato^+^ cells found in the brain parenchyma originated from mature meningeal or perivascular macrophages. To test whether perivascular macrophages can supply the microglial pool in PLX-treated mice, we used *Lyve1^creERT^*^2^*:R26-tdTomato* to transiently label the Lyve1-expressing subset of perivascular macrophages^47,49^ (Fig. S3I-K). After PLX-3x treatment, we did not find any tdTomato^+^ cells in the brain parenchyma of *Lyve1^creERT^*^2^*:R26-tdTomato* mice, suggesting that Lyve1^+^ meningeal macrophages are not capable of infiltrating the brain to supply microglia (Fig. S3L). Therefore, we conclude that brain-engrafted MDMs experience transient expression of CD206, similar to microglia precursors during development.

### Brain-engrafted MDMs share some transcriptional and epigenetic features with meningeal macrophages

Brain-engrafted MDMs likely traverse the meningeal space before entering the brain parenchyma, suggesting that they may acquire transcriptional and epigenetic characteristics similar meningeal macrophages due to environmental imprinting. Alongside the upregulation of *Mrc1*, we observed a substantial overlap of genes upregulated in brain-engrafted MDMs with the signature genes of border-associated macrophages (BAMs), while YS-derived microglia showed minimal similarity to BAMs (Fig. 3F). GSEA confirmed an enriched BAM score in MDMs, indicating shared transcriptional signatures between MDMs and BAMs (Fig. S3M).

Although brain-engrafted MDMs do not exhibit similar protein-level expression of BAM markers such as CD206, epigenetic profiling can uncover the history of transcriptional control and cell state changes. To determine if brain-engrafted MDMs exhibit epigenetic similarities with meningeal macrophages, we performed CUT&TAG for H3K27ac antibody, a histone marker for active promoters and enhancers^52^, on YS-derived microglia, brain-engrafted MDMs, isolated dura macrophages and PBMs (Fig. 3G). Our findings revealed numerous peaks in the promoter regions of meningeal macrophage-associated genes such as *Mrc1* and *Aoah* in MDMs, dura macrophages, and PBMs, while YS-derived microglia showed no peaks in these areas (Fig. 3H). Conversely, YS-derived microglia obtained higher H3K27ac counts at the promoter region of microglia-specific transcription factor Spalt-like transcription factor 1 (*Sall1*) (Fig. 3I). Genes upregulated in mature meningeal macrophages, such as *Lyve1*^47,53^, showed peaks only in dura macrophages and PBMs (Fig. 3J). Thus, we identified some shared transcriptional and epigenetic signatures among dura macrophages, PBMs, and MDMs, while YS-derived microglia remain distinct from these macrophage groups. Our analysis suggests that as MDMs engraft the brain parenchyma and undergo a similar tissue imprinting process as CD206^+^ macrophages in the meningeal space (Fig. S3N).

### Blood-derived cells and skull-derived cells are both capable of replenishing microglia

Lastly, we sought to delineate the anatomical sources of the monocytes that give rise to brain-engrafted MDMs in models of microglial depletion. Studies have shown that circulating Ly6C^high^ CCR2^+^ blood monocytes can engraft the brain if the head is exposed to irradiation or busulfan^10,20,54^. We recently showed that both monocytes and neutrophils residing in the skull bone marrow can directly access the dura mater via skull−dura channels^33,55^, and that in neuroinflammatory conditions these cells can also migrate to the CNS parenchyma^33,56^. Therefore, we wondered whether monocytes from both blood and skull bone marrow can supply brain-engrafted macrophages in microglial depletion model.

To trace blood-derived cells, we established a parabiotic pair by connecting the circulatory systems of ubiquitin-green fluorescent protein (UBC-GFP) and wild-type (WT) mice, enabling donor-derived circulating cells to be identified through GFP expression. Two weeks after surgery, the paired mice underwent either a single (PLX-1x) or repetitive (PLX-3x) treatment (Fig. 4A). In control mice (0.02% ± 0.006) and in mice subjected to the PLX-1x regimen (0.017% ± 0.003), hardly any GFP^+^ donor-derived macrophages were found in the WT host brain (Fig. 4B-D), consistent with previous reports^12^. Following PLX-3x treatment, however, GFP^+^ cells could be observed in the brain parenchyma (6.155% ± 2.054) (Fig. 4B-D). Therefore, we conclude that blood-derived cells can engraft the brain during microglia repopulation.

**Figure 4.**
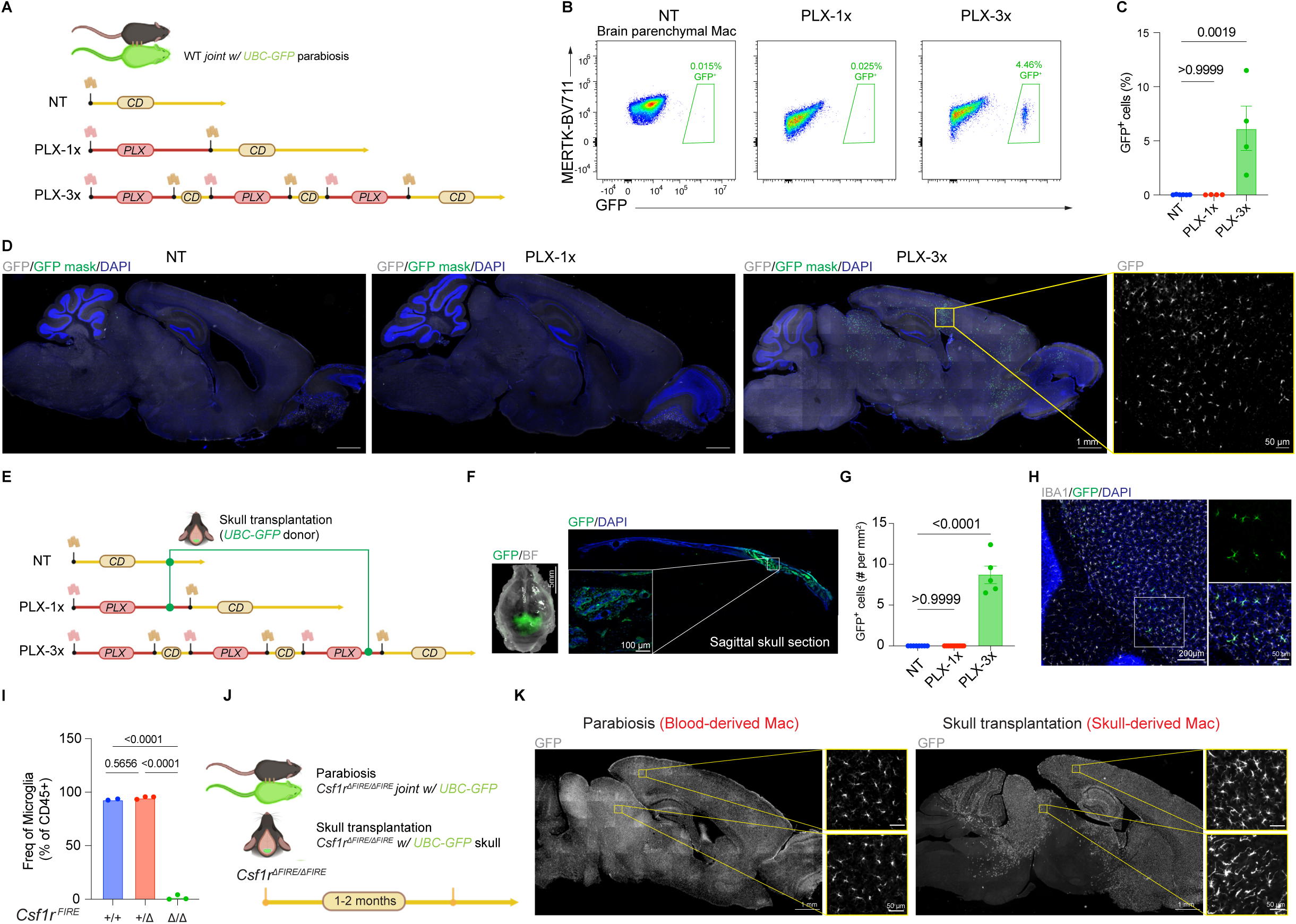
Blood-derived cells and skull-derived cells are both capable of replenishing microglia. (A) Experimental scheme. WT and UBC-GFP donor parabionts received either control diet (NT), PLX-1x, or PLX-3x treatment. (B) Representative flow plot of GFP^+^ brain parenchymal macrophages from different treatment conditions (NT n=6, PLX-1x n=4, PLX-3x n=4). (C) Quantification of the percentage of GFP^+^ brain parenchymal macrophages, one-way ANOVA with Kruskal-Wallis test and Dunn’s multiple comparison test. Data pooled from two independent experiments. (D) Representative IHC images of GFP^+^ cells in sagittal brain sections from different treatment conditions. GFP^+^ cells are masked in green, and the GFP signal is shown in grey, scale = 1 cm. (E) Experimental scheme. WT mice treated with either control diet, PLX-1x, or PLX-3 received skull transplants from UBC-GFP donor mice. (F) Stereotactic image of skull with intact dura, and IHC of imaging of sagittal skull section showing the transplanted GFP^+^ skull tissue in WT mouse, inset shows enlarged region of GFP^+^ transplanted skull tissue. (G) Quantification of GFP^+^ cells per square millimeter in brain parenchyma adjacent to transplant site of mice that received a skull transplant from UBC-GFP donor mice (NT n=8, PLX-1x n=11, PLX-3x n=5). Analyzed with one-way ANOVA with Kruskal-Wallis test and Dunn’s multiple comparison test. (H) Representative IHC image showing GFP^+^ IBA1^+^ cells in brain parenchyma adjacent to the transplant site. (I) Flow cytometry quantification of microglia frequency in *Csf1r^ΔFIRE^*mice, (+/+ WT n=2, +/*Δ* heterozygous n=3, *Δ*/*Δ* homozygous n=3), one-way ANOVA with Tukey’s post-test. (J) Experimental scheme. Homozygous *Csf1r^ΔFIRE/ΔFIRE^* mice underwent either parabiosis or skull transplantation using UBC-GFP donor mice. (K) Immunofluorescent images showing engrafted GFP^+^ cells (in grey) in the brains of homozygous *Csf1r^ΔFIRE/ΔFIRE^* mice after parabiosis (left) or skull transplantation (right).

To determine the potential of skull bone marrow as an alternative source for repopulated microglia^33^ (Fig. 4E-F), we transplanted the 4x6 mm^2^ calvarium bone flap from the UBC-GFP mouse into the WT mouse. Through vascular remodeling, the transplanted GFP^+^ calvarium BM was successfully incorporated into the skull of the WT mouse (Fig. 4E-F), such that all GFP^+^ cells in this chimeric animal model originated only from the transplanted calvarium skull bone. Notably, these GFP^+^ cells were localized to the transplantation area and scarcely appeared in the blood circulation, indicating that skull transplantation is a good model for the specific tracking of skull-derived cells (Fig. 4F and Fig. S4A-B). By means of this technique we administered the two different PLX treatment regimens to WT mice, in each case proceeding with skull transplantation to trace skull-derived cells. Skull-derived cell engraftment in the brain was observed exclusively in mice subjected to repeated PLX-3x treatment (Fig. 4G-H). GFP^+^ cells in the brain also expressed IBA1, confirming their macrophage identity. Co-staining of GFP^+^ cells with endothelial cell marker CD31 revealed the parenchymal localization of GFP^+^ cells (Fig. S4C). Interestingly, although no GFP^+^ parenchymal macrophages were found after PLX-1x, GFP^+^ CD206^+^ macrophages could be observed at the brain border, including those in the leptomeninges and perivascular space (Fig. S4D-E). Furthermore, we employed optical clearing techniques^57^ to delineate the distribution of GFP^+^ cells within the brain after PLX-3x treatment (Movie 1). Interestingly, skull-derived GFP^+^ cells were localized only in the midbrain and cerebellum, where they were positioned proximal to the leptomeninges. This distribution pattern suggests a direct transmeningeal entry of skull-derived cells into the brain. Through the application of skull transplantation, our findings demonstrated that skull-derived cells can contribute to the repopulated microglia pool.

Although PLX5622 is widely used in studies of microglia turnover, pharmacological inhibition of CSF1R might have off-target effects. To determine whether microglial loss alone is sufficient to prompt engraftment of MDMs into the brain, we incorporated a genetic model of microglia deficiency into our study. The *Csf1r* locus contains a highly conserved super-enhancer, known as the fms-intronic regulatory element (*FIRE*), whose deletion in mice selectively depletes certain tissue-resident macrophages such as microglia^58^. Homozygous *Csf1r*^Δ*FIRE/*Δ*FIRE*^ mice have no microglia in the brain (Fig.4I and Fig. S4F) and can thus serve as a good model to establish whether microglial loss is sufficient to trigger blood-monocyte or skull-monocyte engraftment into the brain. To address this question, we cojoined *Csf1r*^Δ*FIRE/*Δ*FIRE*^ mice with UBC-GFP mice to generate parabiotic pairs, while a separate group of *Csf1r*^Δ*FIRE/*Δ*FIRE*^ mice received the skullcaps from UBC-GFP donor mice (Fig. 4J). Two months post-surgery, GFP^+^ cells could be found in the brain parenchyma in both groups of the *Csf1r*^Δ*FIRE/*Δ*FIRE*^ mice (Fig. 4K). In all *Csf1r*^Δ*FIRE/*Δ*FIRE*^ mice joined with UBC-GFP parabionts, GFP^+^ cells were detected in the blood and peripheral organs (Fig. S4G-H); moreover, GFP^+^ cells were evenly distributed across all brain regions, suggesting that these cells enter the brain by crossing the brain vasculature. In contrast to parabiotic pairing, no GFP^+^ cells were detected in the blood or peripheral organs of the *Csf1r*^Δ*FIRE/*Δ*FIRE*^ mice following calvaria transplantation. Intriguingly, skull-derived GFP^+^ macrophages were not distributed across all brain regions (Fig. 4K). In some skull-transplanted mice, GFP^+^ cells were confined to the region beneath the transplantation sites (Fig. S4I-J), suggesting a direct transmigration from skull to dura via dural channels^56,59,60^ and then subsequently from dura to the brain parenchyma. Finally, to determine if brain-blood barrier (BBB) integrity for hydrophilic molecules was compromised in models of microglial depletion, we assessed BBB permeability using sodium fluorescein (NaFl) in *Csf1r*^Δ*FIRE/*Δ*FIRE*^ mice and mice treated with PLX-3x (Fig. S4K). In both cases, our results showed no significant differences in NaFl permeability into the brain between control mice and those lacking microglia, suggesting that MDM engraftment does not require compromised tight junctions (Fig. S4L-M). Therefore, we found that microglial deficiency is sufficient to enable the peripheral cell engraftment into the brain from both blood and skull sources. The diverse origins and potentially different routes of entry may confer unique signatures and functions on blood-derived and skull-derived macrophages.

## DISCUSSION

Here, we compared YS-derived microglia with an under-appreciated population, namely brain-engrafted MDMs, that compose a small subset of parenchymal macrophages in the homeostatic brain and expand substantially upon prolonged microglia depletion. Our findings showed that the ontogeny of brain macrophages dictates their cellular features. The presence of brain-engrafted MDMs has previously been described in pathological conditions such as traumatic brain injury, stroke, and neuroinflammation, as well as in genetic models of microglial defects^21,22^. In alignment with previous studies, we found that although MDMs engraft the brain environment they do not acquire a homeostatic microglial signature that phenocopies YS-derived brain macrophages. Specifically, MDMs do not acquire high expression of SALL1, a critical regulator defining microglial identity and function^61^. Recent studies showed that SALL1 enforces a microglia-specific transcriptional program through its interactions with SMAD4, a transcription factor downstream of transforming growth factor β signaling. Deletion of *Smad4* results in a developmental arrest of microglia, evidenced by the loss of homeostatic microglial signatures and upregulation of with upregulation of PBM genes like *Apoe, Mcr1, Cybb and Ms4a7*^62,63^. This signature is largely consistent with the MDM phenotype described in our study, suggesting that MDMs may transiently differentiate into PBM-like cells at the brain borders, before infiltrating the brain parenchyma. Understanding how ontogeny and tissue imprinting dictate the identity of brain macrophages will require further investigation, potentially through additional epigenetic profiling of YS-derived microglia and brain-engrafted MDMs.

Our study also illuminates the potential of blood-derived and skull-derived cells to replenish microglia, with each source potentially imparting unique characteristics to the brain-engrafted MDMs. Recent studies revealed that cerebrospinal fluid (CSF) can enter the skull bone marrow and influence cranial hematopoiesis^64,65^. This suggests that immune cells in skull bone marrow, upon receiving brain-derived signals from the CSF, may acquire unique properties compared to cells in other bone marrows. Indeed, Kolabas *et al*. found that mouse skull exhibits a unique transcriptomic profile, both in healthy and injured states, characterized by a late-stage neutrophil phenotype^66^. Our studies suggested that brain-engrafted macrophages from the periphery may be heterogeneous, thus a more comprehensive analysis is warranted to study their signatures based on anatomical sources and routes of cell entry.

Lastly, the potential of microglial replacement is becoming increasingly recognized in cell-based therapies for CNS disorders^46,67–69^. Current microglial replacement strategies, however, often require aggressive preconditioning approaches such as irradiation or chemotherapy (e.g. busulfan)^20,46,54^, sometimes combined with intraparenchymal injection of donor cells. Here, we introduce a less aggressive scheme for replacing microglia with peripheral monocytes. We showed that substantial engraftment of MDMs can occur in WT mice after repetitive PLX5622 treatment, without the need for irradiation or myeloablation. Our research suggests that donor cells from both skull bone marrow and blood can infiltrate the CNS, expanding the potential for microglial replacement therapies. Future research could integrate novel microglia replacement strategies with different routes of cell delivery revealed by our study. Deeper understanding of MDM brain infiltration and function will pave the way to targeted therapies that can more effectively modulate the brain’s immune responses, thereby potentially improving outcomes of injury and of neurodegenerative diseases.

## ACKNOWLEDGMENTS

We thank Daniel Gibson and Shirley Smith for editing the manuscript. We thank Mathew Blurton-Jones (University of California, Irvine), Clare Pridans (University of Edinburgh) and David Hume (Mater Research Institute-University of Queensland) for generating and kindly sharing the *Csf1r*^Δ*FIRE/*Δ*FIRE*^ mice used in this study; members of Colonna and Kipnis labs for the valuable suggestions regarding the data acquisition and analysis; Jennifer Ponce from McDonnell Genomic Institute (Washington University in St. Louis) and Amanda Swain (10x Genomics) for the assistance with the single-cell RNA-sequencing experiments; Erica Lantelme, Dorian Brinja and Pascaline Akitani from the flow cytometry core facility at the Pathology and Immunology department; Peter Baguynov from the Center for Cellular Imaging for help and instruction of the tissue clearing and imaging; Abena Apaw and Larysa Kisselbach for taking care of the mice used in this study. The work was supported by grants from NIH (AG078106) and Cure Alzheimer’s Fund to J.K..

## AUTHORS CONTRIBUTIONS

Conceptualization: S.D., M.Co., J.K.; Methodology, S.D., A.D., Y.C., S.S., J.R., T.M., B.B., I.S., S.B.; Software: S.D., P.R., F.O., J.N., J.C.; Validation: S.D.; Formal Analysis: S.D., Y.C., A.D.; Investigation: S.D.; Resources: F.G., M.C., J.K., and M.Ce.; Data curation: S.D.; Writing Manuscript: S.D.; Review & Editing, S.D., D.Q., J.K., and M.Co.; Funding Acquisition: J.K., M.Co.; Supervision: M.Co., and J.K.

## STAR METHODS

### RESOURCE AVAILABILITY

### Further information and requests for resources and reagents should be directed to the lead contact, Jonathan Kipnis

#### Materials availability

Materials and reagents used in this study are listed in the key resources table. This study did not generate new unique reagents.

#### Data and code availability

- Fastq files and quantified gene counts for single-cell sequencing are available at the Gene Expression Omnibus (GEO) with the accession number GSExxxx.
- Cut and tag data for microglia and MDMs (GEO: GSExxxx) and meningeal macrophages (GEO: GSExxxx) are available at the GEO.
- Single-cell RNA-seq for normal microglia and microglia replacement from different transplantation strategies was previously published and reanalyzed in this study (GSE155499)^46^.
- Any additional information required to reanalyze the data reported in this paper is available from the lead contact upon request.

### EXPERIMENTAL MODELS

#### Animals

Wild-type C57BL/6J mice were bred in-house or acquired from the Jackson Laboratory (JAX stock #000664) and housed under specific pathogen-free conditions at the Washington University School of Medicine animal facility. These mice were kept in a controlled environment with a 12-hour light-dark cycle, and provided standard rodent chow and sterilized water ad libitum unless stated otherwise. For animals treated with PLX5622 diet, the CSF1R inhibitor PLX5622 was provided by Plexxikon Inc and was formulated into rodent diet AIN-76A (Research Diets) at a concentration of 1200 mg/kg chow. The chow consumption was monitored daily. Both male and female mice, aged 2 to 6 months, were utilized in experiments, with matching by age and sex within each experimental cohort.

*Ms4a3-Cre* mice^34^ were crossed with *Rosa26-CAG-LSL-tdTomato-WPRE* mice (Ai9; JAX stock #007909)^72^.

*Flt3^Cre^* mice^36,73^ crossed with *Rosa26-CAG-LSL-EYFP* mice (Ai2; JAX stock # 007920)^72^ bred in house were kindly provided by Dr. David DeNardo.

*Mrc1^creERT^*^2^ and *Lyve1^creERT^*^2^ were generated by CRISPR-Cas9 technology (Cyagen). gRNA targeting mouse Mrc1 gene (AGAAGCCTCATGGCCCAGAGTGG), or Lyve1 gene (GAAGTTTAGATGCAAGAGAGTGG) was co-injected with the donor vector containing “CreERT2-P2A” cassette, and Cas9 mRNA into fertilized mouse eggs to generate targeted knockin offspring. F0 founder animals were identified by PCR followed by sequence analysis, which were bred to wildtype mice to test germline transmission and F1 animal generation.

For genotyping of the *Mrc1^creERT^*^2^ mice the following primers were used: Fwd. TCCTCTTCCTCCGTTACCTGAAA; Rev. CCAAGTGGCTTTGGTCCGTC. For genotyping of the *Lyve1^creERT^*^2^ mice the following primers were used: Fwd. TTCATCCCCTGACTCCACAACA; Rev. CATAGAGGGGCACCACGTTCT.

All experimental procedures involving animals were conducted following protocols approved by the Institutional Animal Care and Use Committee at Washington University in St. Louis (protocol #21-0145).

#### Sample preparation for flow cytometry and FACS

<u>For brain,</u> sample preparation was executed on ice to preserve tissue integrity. Mice were administered a lethal dose of Euthasol (10% v/v) intraperitoneally, followed by transcardial perfusion with PBS containing 0.025% heparin. Post-decapitation, brains were excised and immediately submerged in cold RPMI (Gibco). For FACS, the choroid plexus was carefully extracted and discarded. Brain tissue was then dissociated using a dounce homogenizer, filtered through a 70 μm cell strainer into a 50 ml conical tube, and centrifuged for 10 mins/450xg at 4°C. The resulting brain cell pellet was resuspended in a 30% isotonic percoll solution (GE Healthcare) to remove myelin, and myelin fraction was depleted by centrifugation for 15 mins/850xg at 4°C (acceleration 6, break 1). The clean cell pellets were then resuspended in a FACS buffer containing 1% BSA (Equitech-Bio), 1mM EDTA (Corning), and 10mM HEPES (Corning) in preparation for staining.

<u>For dura</u>, skull caps were collected into ice-cold RPMI (Gibco). Meninges were then peeled from the skull cap using fine forceps Tissues underwent enzymatic digestion in a pre-warmed digestion buffer of DMEM with 2% FBS, 1 mg/mL collagenase VIII (Sigma Aldrich), and 0.5 mg/mL DNase I (Sigma) at 37°C with agitation for 15 mins. Post-digestion, the tissues were filtered through a 70 μm cell strainer, and digestion was halted with an equal volume of complete medium (DMEM with 10% FBS). The samples were then diluted with FACS buffer, centrifuged at 450xg for 5 minutes, and the pellet resuspended in FACS buffer, kept on ice until further use.

<u>For blood,</u> sample was collected into 1 mL of ice-cold PBS with 20mM EDTA to prevent coagulation. Blood was centrifuged at 450xg for 10 minutes, and red blood cell (RBC) lysis was performed by resuspension in 1 mL of ACK lysis buffer (Quality Biological) for 5 minutes, and then washed with PBS. Following a final centrifugation for 10 mins/450xg, the supernatant was discarded, leaving the leukocyte-rich pellet for subsequent analysis.

#### Flow cytometry analysis

Prior to staining, Fc-receptor blockade was performed using anti-CD16/32 (Biolegend) diluted 1:50 in FACS buffer. The sample was incubated for 10 min on ice. Next, conjugated antibodies were added for 20 minutes at 4°C. Flow cytometry analysis was performed Cytek Aurora spectral flow cytometer (Cytek) then analyzed using FlowJo software (Tree Star).

#### Fluorescence-activated cell sorting (FACS)

For sorting experiments, cells underwent Fc-receptor blockade as above, followed by surface staining for 25 min on ice. To sort brain parenchymal macrophages for bulk RNA-seq experiment, anti-CD11b, anti-CD45 and lineage antibody cocktail (anti-Ly6C, anti-Ly6G, anti-CD206, anti-CD44) were used. Targeted cells were sorted into cold RLT Plus buffer (from RNeasy Micro Plus Kit, RQiagen) supplied with 2-Mercaptoethanol (Sigma). To sort brain immune cells for scRNA-seq experiment, TotalSeq-B hashtag antibodies for CD45 and MHC, anti-CD45.2 and anti-CD11b were used. After staining, cells were incubated in FACS buffer supplied with 0.2 μg/mL DAPI (Sigma). Sorting was performed on FACS Aria-II (BD Bioscience). Cells negative for DAPI were selected for being live. Targeted cells were sorted into sterile PBS with 2% BSA (Equitech-Bio), then were washed with 0.04% ultrapure BSA (Thermo Fisher) in PBS before loading to the chip.

#### Bulk RNA-seq

Brain parenchymal macrophages were purified from PLX5622-treated mouse brains using FACSAria II (BD Biosciences) as live, negative for Ly6C, Ly6G, CD206, CD44; positive for CD45 and CD11b. From the brain parenchymal macrophage gate, tdTomato^+^ and tdTomato^-^ macrophages were sorted into different tubes containing cold RLT Plus buffer (from RNeasy Micro Plus Kit, RQiagen) supplied with 2-Mercaptoethanol (Sigma). Next, total RNA was extracted from samples using an RNeasy Micro Plus Kit (Qiagen). Total RNA integrity was determined using Agilent Bioanalyzer or 4200 TapeStation. Double standard complementary DNA (cDNA) was prepared using the SMARTer Ultra Low RNA kit for Illumina Sequencing (Takara-Clontech) per manufacturer’s protocol. cDNA was fragmented using a Covaris E220 sonicator using peak incident power 18, duty factor 20%, and cycles per burst 50 for 120 s. cDNA was blunt ended, had an A base added to the 3′ ends, and then had Illumina sequencing adapters ligated to the ends. Ligated fragments were then amplified for 12 to 15 cycles using primers, incorporating unique dual index tags. Fragments were sequenced on an Illumina NovaSeq-6000 using paired end reads extending 150 bases. RNA-seq reads were then aligned and quantitated to the Ensembl release 101 primary assembly with an Illumina DRAGEN Bio-IT on-premise server running version 3.9.3-8 software.

Aligned gene counts were processed using the DESeq2 package with R. Genes with fewer than 10 counts among all of the samples were excluded. DEGs were defined as protein-coding genes with an average expression > 100 counts and false discovery rate < 0.05. Gene Set Enrichment Analysis of hallmark genes was performed using fgsea (Korotkevich G, Sukhov V, Sergushichev A (2019)).

#### Parabiosis

*UBC-GFP* mice, together with sex and age matched WT mice were rendered fully muscle-relaxed via intraperitoneal injection of a xylazine (10 mg/kg) and ketamine (100 mg/kg) mixture. The respective lateral regions were first shaved, then applied with betadine and alcohol in three alternating cycles. The surgery was performed using aseptic technique. Next, a cut extending from the olecranon to the knee joint was made, allowing for the separation of the subcutaneous fascia to yield a 0.5 cm margin of free skin. The olecranon and knee joints were then conjoined with 5-0 Vicryl absorbable sutures. The dermis of the parabiotic pairs was approximated, deliberately excluding epidermal layers at the junction, and sutured closed. Postoperative care included Buprenorphine SR (1.0 mg/kg) and antibiotic administration in drinking water.

#### Calvarium bone-flap transplantation

Mice were anesthetized using a xylazine (10 mg/kg) and ketamine (100 mg/kg) mixture. Throughout the surgical procedure, body temperature was maintained at 37°C using a heating pad. For the initial preparation, the heads of WT mice were shaved and a midline incision was made to expose the skull. A cranial window (4-mm-by-6-mm) was drilled into the parietal and interparietal bones of the skull using an electric drill. To prevent thermal damage, the drilling site was intermittently cooled with 0.9% sodium chloride solution. After the cranial window was detached, careful craniotomy was conducted without damaging the underlying dura mater, brain, or sinus structures. Simultaneously, a sex-matched UBC-GFP mouse was used as a skull BM donor. After euthanization and decapitation, a piece of the donor’s calvarium, matching the host’s window size, was harvested. The periosteum and pericranium were left intact on the bone flap to enhance survival post-transplant. The meninges were removed from the donor skull piece containing the bone marrow, which was then placed within the surgical window of the host. The donor skull was affixed to the host site using cyanoacrylate glue and secured with a 10-0 nylon suture. Post-surgery, the incision was closed, and mice were kept on a heating pad until recovery. Postoperative care included Buprenorphine SR (1.0 mg/kg) and antibiotic administration in drinking water for the first five days to mitigate inflammation and prevent infection.

#### Immunohistochemistry

After perfusion, tissues were preserved in 4% paraformaldehyde (Electron Microscopy Sciences) at 4°C for an overnight period. Subsequently, skull bones were decalcified in 0.5M EDTA (Corning) for 72 hours. The fixed tissues were then subjected to a dehydration process in a 30% sucrose solution until they submerged fully, followed by encapsulation in Tissue-Plus OCT compound (Thermo Scientific) and storage in a freezer. During sectioning, the cryo-embedded blocks containing either brain or skull tissues were sectioned into 40μm-thick slices using a cryostat (Leica). Staining of the free-floating sections began with a 2-hour block in PBS with 5% BSA and 0.5% Triton X-100. Primary antibodies were applied in PBS with 1% BSA and 0.5% Triton X-100 and left to bind overnight at 4°C. This was followed by secondary antibody staining with fluorochrome conjugates at room temperature for 2 hours. DAPI (Sigma) or TO-PRO-3 iodide (Thermo Fisher Scientific) was added to the secondary antibody mixture to counterstain for nuclei. Between each staining step, sections were thoroughly rinsed with 0.5% Triton X-100 in PBS. Lastly, the stained sections were mounted on Superfrost glass slides (Fisher Scientific) and sealed with mounted with FluorSave^TM^ (Millipore) mounting media.

#### Confocal and widefield microscopy

Brain, dura or skull sections were acquired using a wide-field fluorescent microscope (Lecia). Quantitative analysis of imaging measurements was performed using the FIJI package for ImageJ. For coverage measurements, images were thresholded to identify positive signal, and were quantified as area of signal/total area of site. All imaging analysis and threshold settings were done blindly, to allow for no experimental bias. For cell counting in skull transplantation experiment, a 1x1 mm^2^ square was drawn from the top of brain underneath the transplanted area as area of interest. GFP^+^ cells per area of interest were recorded. At least five images per animal were averaged to generate the value utilized for a single mouse. The cell counting was done blindly to avoid experimental bias. For sholl analysis, at least 15 cells per animal were chosen randomly to generate the averaged value for a single mouse.

Confocal imaging of brain cryosections was performed using a TCS SP8 inverted confocal microscope. 20 μm-thick z-stack images were acquired with a 20X air objective at 1024×1024-pixel or 2048x2048 resolution, z-step=1μm, line averaging=2. Maximal projections were rendered in Imaris. The measurement of CD68 volume was determined with Imaris via its Surface function. Positive signals for CD68 within z stacks were used to create Surfaces. Identical setting was applied to create Surface for a set of images with the same staining.

#### Tissue clearing and lightsheet microscopy

Brains were collected and immediately fixed in 4% paraformaldehyde (Electron Microscopy Sciences) and maintained at 4°C for a duration of one day. For staining, Fluotag-X4 anti-GFP sdAb AF647 (NanoTag Biotechnologies) was prepared in a dilution of 1:200 in PBS containing 0.5% Triton X-100. The brains were then immersed in this antibody solution and incubated at 4°C with gentle agitation for one week. Following incubation, the brains were washed thrice in PBS, each for 5 minutes. A series of methanol dehydration steps followed, involving 20%, 40%, 60%, and 80% methanol solutions in PBS for a minimum of one hour each, and then two sequential overnight incubations in 100% methanol. Subsequent to methanol dehydration, the heads were processed with an overnight incubation in a 2:1 mixture of dichloromethane to methanol, and then in 100% dichloromethane for another night. For optical clearing, heads were submerged in dibenzyl ether overnight at room temperature. The final step involved transferring the specimens to ethyl cinnamate for refractive index matching, also overnight. The cleared brains were imaged in a bath of ethyl cinnamate using an UltraMicroscope II light sheet fluorescence microscope (Miltenyi Biotec).

#### Sodium Fluorescein Permeability Assay

Sodium Fluorescein salt was dissolved in sterile PBS to a final concentration of 5 mg/ml. Mice were injected with 50 mg/kg Sodium fluorescein solution intraperitoneally. After 1 hour, blood was sampled and mice were then perfused with D-PBS for 3 minutes. Brains were dissected and one hemisphere was frozen on dry ice. The blood samples were then centrifuged at 100xg for 10 min and the supernatant (plasma) was collected and plasma and brain were both stored at −80°C until homogenization. Brain tissue was homogenized in 7 volumes of PBS at max. power for 90 sec using a Fisherbrand™ Bead Mill 4 Mini Homogenizer. The brain homogenates were then centrifuged, and the resulting supernatant was diluted 1:1 in 2% TCA. Plasma samples were diluted 1:140 in sterile PBS, followed by an additional 1:1 dilution in 2% Trichloroacetic Acid (TCA). Brain and plasma samples were incubated overnight rotating at 4°C. Both sets of samples were then centrifuged and the supernatants were diluted 1:1 in borate buffer, pH 11. The samples were then added to a 96-well plate with 3 technical replicates per sample. The plate was analyzed on a BioTek Synergy H1 plate reader (Excitation=480nm, Emission=538nm).

#### Single-cell RNAseq analysis

The Seurat (v4.0) package in R was used for analysis. Filtered gene-by-cell matrices of UMI counts for public datasets were read into R using the Seurat Read10X function and converted into S4 objects using the CreateSeuratObject function from the same package. For quality control, cells with low UMI and gene number per cell were filtered out. Cutoffs for UMI and gene number were empirically determined on the basis of histograms showing cell density as a function of UMI per gene counts. Genes expressed in fewer than 10 cells were removed from the dataset. Expression values were then normalized using the Seurat’s NormalizeData function. The integrated filtered and normalized matrix was used as input to the Seurat v4 pipeline^74^ and cells were scaled across each gene. These were then integrated using harmony^75^. Gene signature scoring was determined using AddModuleScore function^76^.

### QUANTIFICATION AND STATISTICAL ANALYSIS

All statistical analyses display mean value ± standard error of the mean (SEM). Statistical significance was set at p-value <0.05. GraphPad Prism software was utilized for statistical analysis. A p-value < 0.05 indicated significant differences between conditions, ascertained through various tests: a two-tailed unpaired t-test, parametric paired t-test, ordinary one-way ANOVA, one-way ANOVA with Dunnett’s multiple comparisons or two-way ANOVA with Sidak’s multiple comparisons, as specified in figure legends.

