## Supplementary Figures for "Brain-Engrafted Monocyte-derived Macrophages from Blood and Skull-Bone Marrow Exhibit Distinct Identities from Microglia"

Sup.Fig.1

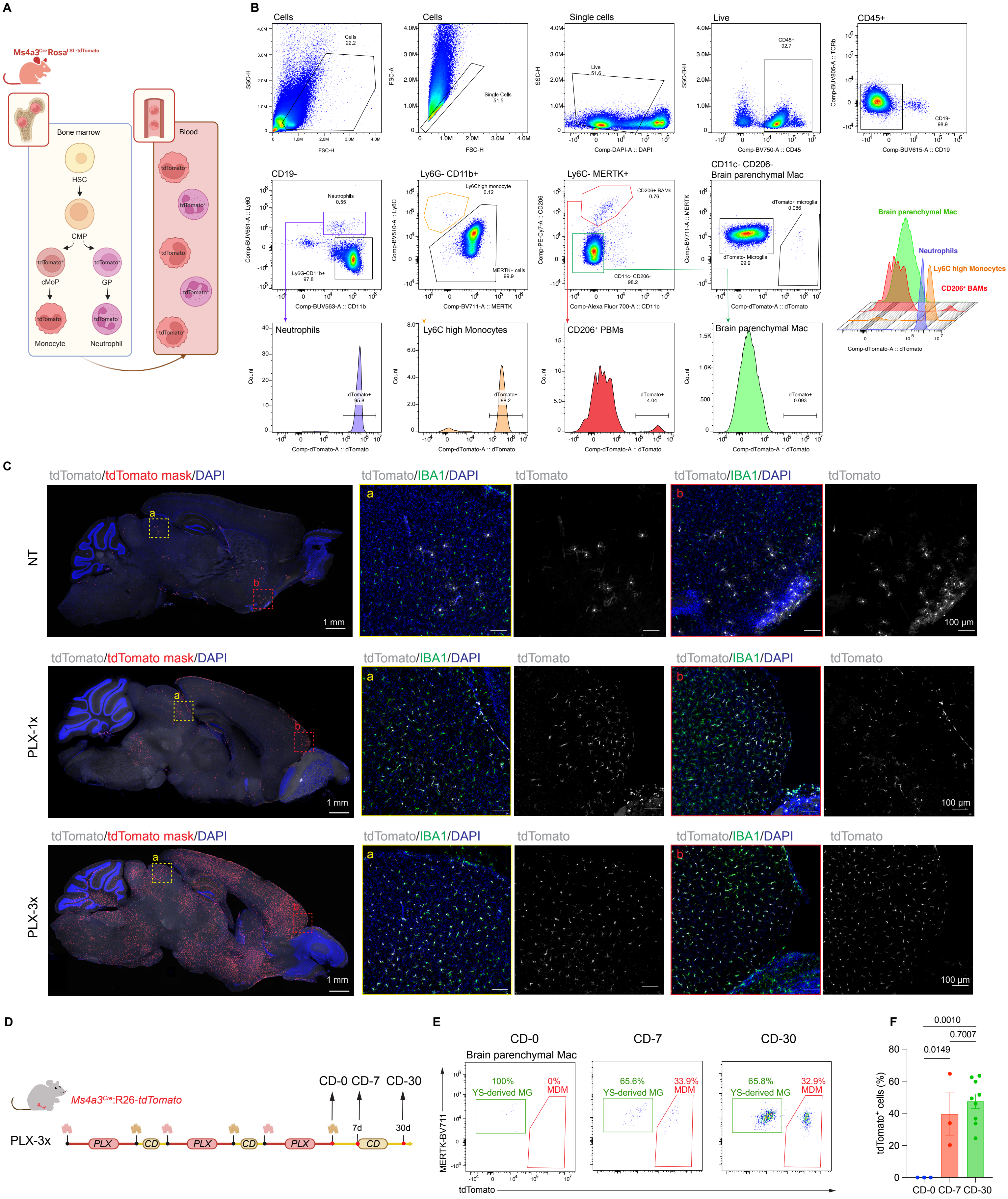
A
B
C
D
E
F
Bone marrow
Blood
HSC
CMP
cMoP
GP
Monocyte
Neutrophil
Cells
Single cells
Live
CD45+
CD19-
Ly6G- CD11b+
Ly6C- MERTK+
CD11c- CD206-
Brain parenchymal Mac
Neutrophils
Ly6C high Monocytes
CD206+ BAMs
Brain parenchymal Mac
Neutrophils
Ly6C high Monocytes
CD206+ BAMs
tdTomato/tdTomato mask/DAPI
tdTomato/IBA1/DAPI
tdTomato
NT
PLX-1x
PLX-3x
1 mm
100 μm
Ms4a3Cre:R26-tdTomato
PLX-3x
PLX
CD
CD-0 CD-7 CD-30
7d
30d
CD-0
CD-7
CD-30
Brain parenchymal Mac
100%
YS-derived MG
0%
MDM
65.6%
YS-derived MG
33.9%
MDM
65.8%
YS-derived MG
32.9%
MDM
MERTK-BV711
tdTomato
0.0010
0.7007
0.0149
tdTomato+ cells (%)
CD-0 CD-7 CD-30

**Figure S1, related to Figure 1**

(A) Cartoon scheme showing the tdTomato expression of *Ms4a3$^{Cre}$:R26-tdTomato* mice in the bone marrow and blood. The graph was made based on Liu et al., 2019[34].

(B) Representative gating strategy used in identifying neutrophils, monocytes, PBMs and brain parenchymal macrophages (including microglia and brain engrafted MDMs). Lower and right panels show the tdTomato expression across different brain immune cells in *Ms4a3$^{Cre}$:R26-tdTomato* mice.

(C) Left panel: Same representative immunofluorescent images showing tdTomato$^{+}$ cells in the brain of NT, PLX-1x, PLX-3x treated *Ms4a3$^{Cre}$:R26-tdTomato* mice as Fig. 1D. Right panel: enlarged immunofluorescent images showing IBA1 (green) and tdTomato (grey) expression in selected regions. scale=100 μm.

(D) Experimental scheme. *Ms4a3$^{Cre}$:R26-tdTomato* mice were treated with PLX-3x, and the brains were harvested 0 day, 7 days or 30 days after control diet (CD) treatment.

(E) Representative flow plot of tdTomato$^{+}$ brain parenchymal macrophages at different time points.

(F) Quantification of the percentage of tdTomato$^{+}$ brain parenchymal macrophages, ordinary one-way ANOVA (CD-0 N=3, CD-7 N=3, CD-30 N=9).

**Sup.Fig2**

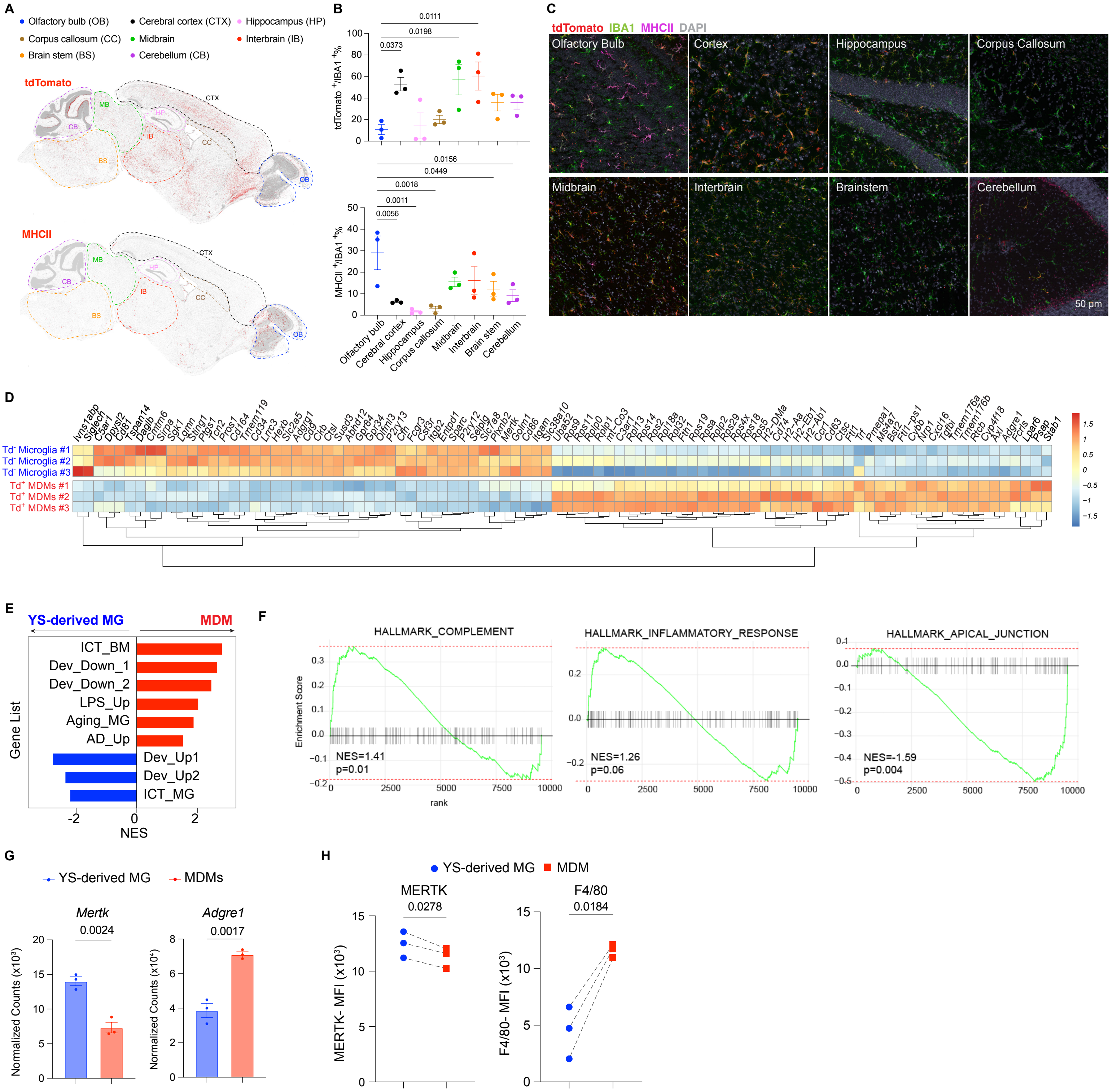
A
Olfactory bulb (OB)
Cerebral cortex (CTX)
Hippocampus (HP)
Corpus callosum (CC)
Midbrain
Interbrain (IB)
Brain stem (BS)
Cerebellum (CB)
tdTomato
MHCII
CTX
MB
HP
CB
IB
CC
BS
OB
B
tdTomato +/IBA1 +%
MHCII +/IBA1 +%
0.0111
0.0198
0.0373
0.0156
0.0449
0.0018
0.0011
0.0056
Olfactory bulb
Cerebral cortex
Hippocampus
Corpus callosum
Midbrain
Interbrain
Brain stem
Cerebellum
C
tdTomato IBA1 MHCII DAPI
Olfactory Bulb
Cortex
Hippocampus
Corpus Callosum
Midbrain
Interbrain
Brainstem
Cerebellum
50 µm
D
Ivns1abp Siglech C5ar1 Dpysl2 Cd81 Tspan14 Daglb Cmtm6 Sirpa Sgk1 Lgmn Sting1 Ptgs1 Tcn2 Pros1 Cd164 Tmem119 Cd34 Lrrc3 Hexb Slc2a5 Adgrg1 Cd9 Ckb Ctsl Susd3 Abhd12 Gpr84 Gpr34 Olfml3 P2ry13 Cfh Fcgr3 Csf3r Itgb2 Entpd1 Sparc P2ry12 Selplg Slc7a8 Plxnb2 Mertk Golm1 Cd86 Itgam Slc38a10 Uba52 Rps9 Rps11 Rplp0 Rplp1 mt-Co3 C3ar1 Rpl13 Rps14 Rps2 Rpl18a Rpl32 Fth1 Rps19 Rpsa Rplp2 Rps29 Rps4x Rps18 Rps5 H2-DMa Cd74 H2-Aa H2-Eb1 H2-Ab1 Ccr1 Cd63 Ctsc Ftl1 Trf Pmepa1 Ms4a7 Bst2 Ftl1-ps1 Cybb Nrp1 Cxcl16 Tgfbi Tmem176a Tmem176b Rtcb Cyp4f18 Axl Adgre1 Fcrls Lpar6 Psap Stab1
Td- Microglia #1
Td- Microglia #2
Td- Microglia #3
Td+ MDMs #1
Td+ MDMs #2
Td+ MDMs #3
1.5
1
0.5
0
−0.5
−1
−1.5
E
YS-derived MG
MDM
Gene List
ICT_BM
Dev_Down_1
Dev_Down_2
LPS_Up
Aging_MG
AD_Up
Dev_Up1
Dev_Up2
ICT_MG
NES
F
HALLMARK_COMPLEMENT
Enrichment Score
NES=1.41
p=0.01
rank
HALLMARK_INFLAMMATORY_RESPONSE
NES=1.26
p=0.06
HALLMARK_APICAL_JUNCTION
NES=-1.59
p=0.004
G
YS-derived MG
MDMs
Mertk
0.0024
Adgre1
0.0017
Normalized Counts (x10³)
Normalized Counts (x10⁴)
H
YS-derived MG
MDM
MERTK
0.0278
F4/80
0.0184
MERTK- MFI (x10³)
F4/80- MFI (x10³)

**Figure S2, related to Figure 1-2**

(A) Representative immunofluorescent image of tdTomato (upper panel) and MHCII (lower panel) expression in the brain of PLX-3x treated *Ms4a3$^{Cre}$:R26-tdTomato* mice. Different brain regions were bounded by the dashed lines. CTX: cortex; HP: hippocampus; CC: corpus callosum; IB: interbrain; MB: midbrain; BS: brainstem; CB: cerebellum.

(B) Quantification of the percentage of tdTomato$^{+}$ cells (upper panel) and MHCII$^{+}$ (lower panel) cells over all IBA1$^{+}$ macrophages across different brain regions (N=3 mice, 3 brain sections per mouse). Analyzed with ordinary one-way ANOVA with Sidak's multiple comparisons test.

(C) Representative confocal image of tdTomato, IBA1, and MHCII staining across different brain regions of PLX-3x treated *Ms4a3$^{Cre}$:R26-tdTomato* mice. Scale = 50µm.

(D) Heatmap showing the differentially upregulated genes in Td$^{-}$ microglia or Td$^{+}$ MDMs.

(E) Normalized enrichment scores (NESs) from GSEA comparing YS-derived MG and MDMs for enrichment in genes upregulated in bone marrow cells (ICT_BM) or microglia (ICT_MG) engrafted to the *Csf1r$^{-/-}$* brain through ICT, AD (AD_Up), after LPS treatment (LPS_Up), changed in during development (DEV_Up 1, 2 or DEV DOWN1, 2). Detailed gene lists are included in Table S1.

(F) GSEA using Hallmark genesets comparing YS-derived MG to MDMs for pathway enrichment.

(G) Normalized RNA counts for Mertk and Adgre1, analyzed with unpaired t-test.

(H) MFI of Mertk expression on PLX-3x treated *Flt3$^{Cre}$:R26-YFP* mice, paired t-test.

Sup.Fig3

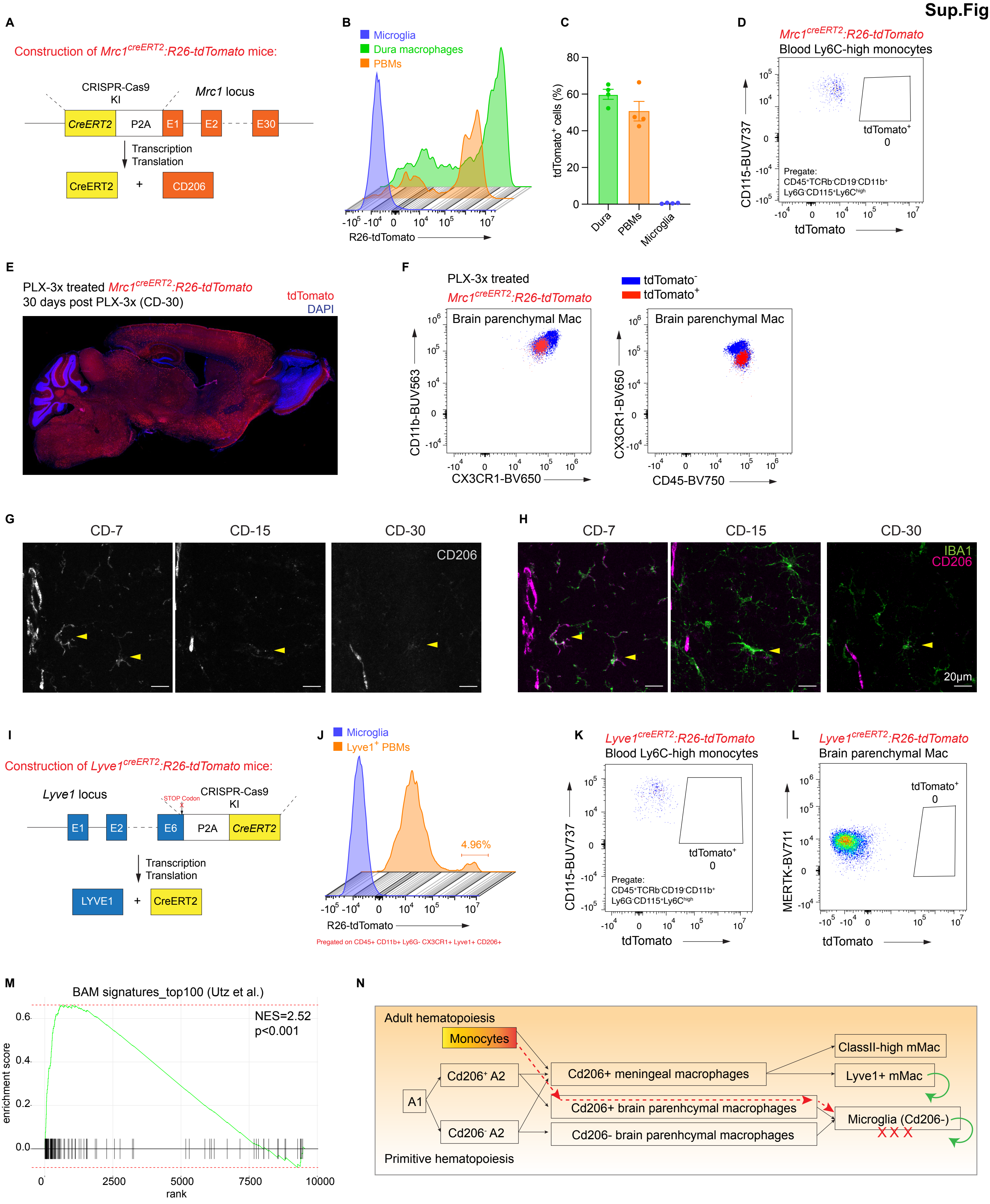
A
Construction of Mrc1creERT2:R26-tdTomato mice:
CRISPR-Cas9 KI
Mrc1 locus
CreERT2
P2A
E1
E2
E30
Transcription
Translation
CreERT2
+
CD206
B
Microglia
Dura macrophages
PBMs
R26-tdTomato
C
tdTomato+ cells (%)
Dura
PBMs
Microglia
D
Mrc1creERT2:R26-tdTomato
Blood Ly6C-high monocytes
CD115-BUV737
tdTomato+
0
Pregate:
CD45+TCRb-CD19-CD11b+
Ly6G-CD115+Ly6Chigh
tdTomato
E
PLX-3x treated Mrc1creERT2:R26-tdTomato
30 days post PLX-3x (CD-30)
tdTomato
DAPI
F
PLX-3x treated
Mrc1creERT2:R26-tdTomato
tdTomato-
tdTomato+
Brain parenchymal Mac
CD11b-BUV563
CX3CR1-BV650
Brain parenchymal Mac
CX3CR1-BV650
CD45-BV750
G
CD-7
CD-15
CD-30
CD206
H
CD-7
CD-15
CD-30
IBA1
CD206
20μm
I
Construction of Lyve1creERT2:R26-tdTomato mice:
Lyve1 locus
STOP Codon
CRISPR-Cas9 KI
E1
E2
E6
P2A
CreERT2
Transcription
Translation
LYVE1
+
CreERT2
J
Microglia
Lyve1+ PBMs
4.96%
R26-tdTomato
Pregated on CD45+ CD11b+ Ly6G- CX3CR1+ Lyve1+ CD206+
K
Lyve1creERT2:R26-tdTomato
Blood Ly6C-high monocytes
CD115-BUV737
tdTomato+
0
Pregate:
CD45+TCRb-CD19-CD11b+
Ly6G-CD115+Ly6Chigh
tdTomato
L
Lyve1creERT2:R26-tdTomato
Brain parenchymal Mac
tdTomato+
0
MERTK-BV711
tdTomato
M
BAM signatures_top100 (Utz et al.)
NES=2.52
p<0.001
enrichment score
rank
N
Adult hematopoiesis
Monocytes
ClassII-high mMac
Cd206+ A2
Cd206+ meningeal macrophages
Lyve1+ mMac
A1
Cd206+ brain parenhcymal macrophages
Microglia (Cd206-)
XXX
Cd206- A2
Cd206- brain parenhcymal macrophages
Primitive hematopoiesis

**Figure S3, related to Figure 3**

(A) Cartoon describing the generation of $Mrc1^{creERT2}$*:R26-tdTomato* mice.

(B) Representative tdTomato expression in microglia, dura macrophages and PBMs of control $Mrc1^{creERT2}$*:R26-tdTomato* mice treated with tamoxifen.

(C) Percentage of microglia (gating strategy in Fig. S1, brain parenchymal macrophage gate), dura macrophages and PBMs (gating strategy in Fig. S1, CD206+ macrophages) expressing tdTomato in control $Mrc1^{creERT2}$*:R26-tdTomato* mice. N=4 mice, single experiment.

(D) Representative flow plot showing tdTomato expression in Ly6C-high monocytes in the blood of $Mrc1^{creERT2}$*:R26-tdTomato* mice treated with tamoxifen.

(E) Representative immunofluorescent image showing tdTomato expression (in red) in the brain of $Mrc1^{creERT2}$*:R26-tdTomato* mice treated with PLX-3x and tamoxifen.

(F) Representative flow cytometry plots of CX3CR1 and CD11b expression (left panel), or CD45 and CX3CR1 expression (right panel) of $tdTomato^-$ (blue) or $tdTomato^+$ (red) brain parenchymal macrophages of the $Mrc1^{creERT2}$*:R26-tdTomato* mice with PLX-3x treatment.

(G) Representative confocal image of CD206 expression (in grey) in the dentate gyrus of mice treated with PLX-3x, following by 7 days (CD-7), 15 days (CD-15) and 30 days (CD-30) of common diet treatment. Scale=20 μm.

(H) Representative confocal image of IBA1 (in green) and CD206 (in magenta) expression in the dentate gyrus of mice treated with PLX-3x, following by 7 days (CD-7), 15 days (CD-15) and 30 days (CD-30) of common diet treatment. Scale=20 μm.

(I) Cartoon scheme describing the generation of $Lyve1^{creERT2}$*:R26-tdTomato* mice.

(J) Representative tdTomato expression in microglia, Lyve1$^+$ PBMs (Pre-gate: CD45$^+$CD11b$^+$ Ly6G$^-$CX3CR1$^+$CD206$^+$Lyve1$^+$) of *Lyve1$^{creERT2}$:R26-tdTomato* mice treated with tamoxifen.

(K) Representative flow plot showing tdTomato expression in Ly6C-high monocytes in the blood of *Lyve1$^{creERT2}$:R26-tdTomato* mice treated with tamoxifen. N=4, single experiment.

(L) Representative flow plot showing tdTomato expression in brain parenchymal macrophages of *Lyve1$^{creERT2}$:R26-tdTomato* mice treated with tamoxifen. N=4, single experiment.

(M) GSEA comparing YS-derived MG and MDMs for enrichment in the top 100 genes upregulated in BAMs compared to microglia from Utz et al.

(N) Cartoon summarizing the dynamic expression of Cd206 in meningeal macrophages and brain parenchymal macrophages during development, and in conditions of depletion (shown by the red crosses). Green circles indicate self-renewal. Dash lines in red suggest the trajectory of brain engrafted MDMs from monocytes.

Sup.Fig4

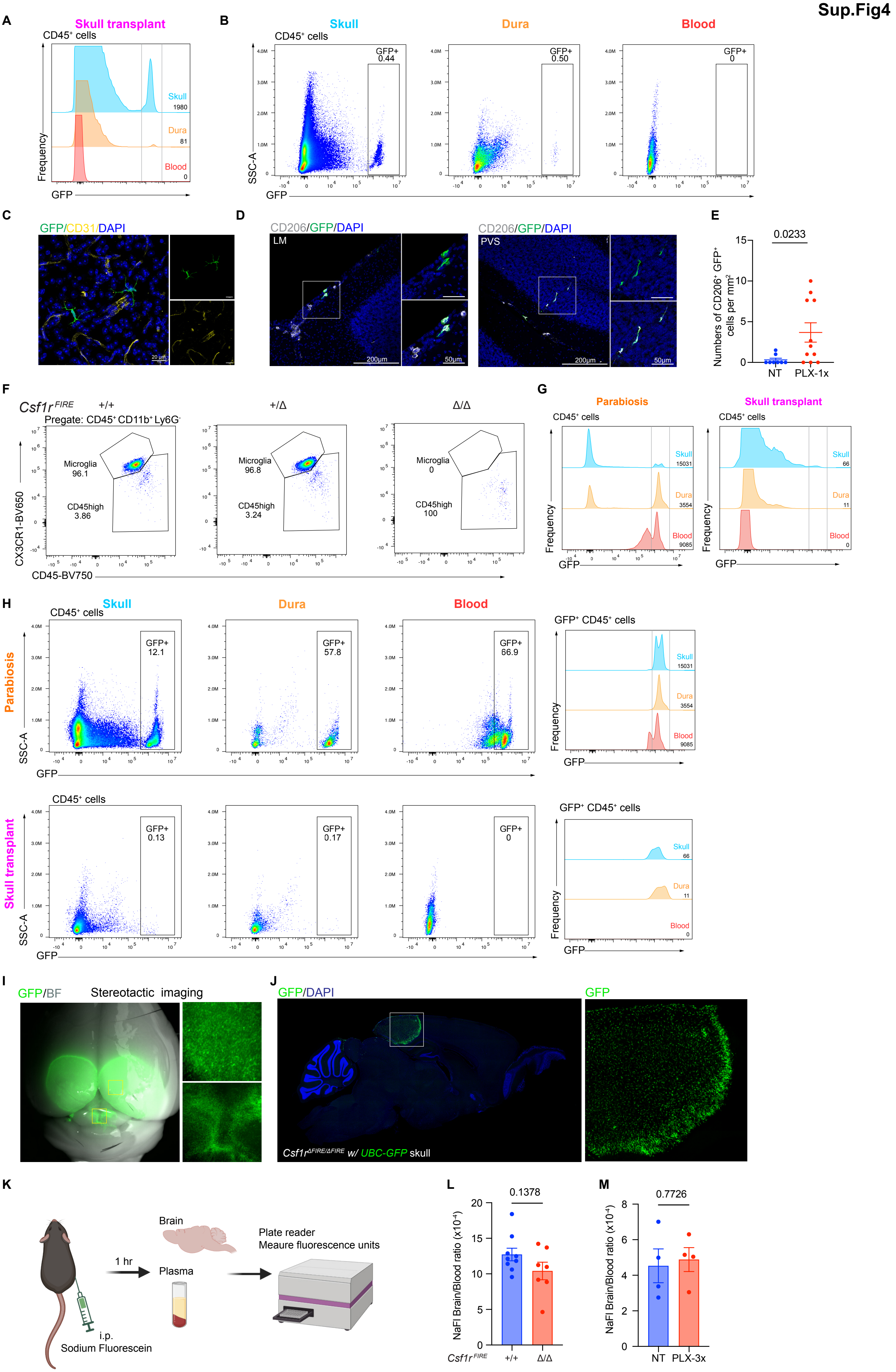

A
Skull transplant
CD45+ cells
Skull
Dura
Blood
Frequency
GFP
B
Skull
Dura
Blood
CD45+ cells
GFP+ 0.44
GFP+ 0.50
GFP+ 0
SSC-A
GFP
C
GFP/CD31/DAPI
D
CD206/GFP/DAPI
LM
PVS
200µm
50µm
E
Numbers of CD206+ GFP+ cells per mm2
0.0233
NT
PLX-1x
F
Csf1r FIRE +/+
+/Δ
Δ/Δ
Pregate: CD45+ CD11b+ Ly6G-
Microglia 96.1
CD45high 3.86
Microglia 96.8
CD45high 3.24
Microglia 0
CD45high 100
CX3CR1-BV650
CD45-BV750
G
Parabiosis
Skull transplant
CD45+ cells
Frequency
GFP
H
Skull
Dura
Blood
Parabiosis
Skull transplant
CD45+ cells
GFP+ 12.1
GFP+ 57.8
GFP+ 66.9
GFP+ 0.13
GFP+ 0.17
GFP+ 0
GFP+ CD45+ cells
SSC-A
GFP
Frequency
I
GFP/BF
Stereotactic imaging
J
GFP/DAPI
GFP
Csf1r ΔFIRE/ΔFIRE w/ UBC-GFP skull
K
Brain
Plasma
1 hr
i.p.
Sodium Fluorescein
Plate reader
Meaure fluorescence units
L
NaFl Brain/Blood ratio (x10-4)
0.1378
Csf1r FIRE
+/+
Δ/Δ
M
NaFl Brain/Blood ratio (x10-4)
0.7726
NT
PLX-3x

**Figure S4, related to Figure 4**

(A) Percentage of GFP$^+$ cells in skull (blue), dura (orange) and blood (red) of WT mice received skull transplantation from UBC-GFP mice. The number indicated the number of cells recorded in the GFP$^+$ gate.

(B) Representative flow plots of GFP$^+$ expression of all CD45$^+$ cells in skull, dura and blood of WT mice received skull transplantation from UBC-GFP mice.

(C) Representative immunofluorescent images showing CD31$^+$ and GFP$^+$ cells in the brain of PLX-3x treated WT mice after skull transplantation (donor: UBC-GFP mice) . scale=20 µm.

(D) Representative confocal images showing the presence of CD206$^+$ GFP$^+$ cells in the leptomeninges (LM) and perivascular space (PVS) of PLX-1x treated WT mice after receiving skull from UBC-GFP mice. scale=200 µm.

(E) Quantification of CD206$^+$ GFP$^+$ cells per square millimeter in leptomeninges adjacent to transplant site of mice that received a skull transplant from UBC-GFP donor mice (NT n=8, PLX-1x n=11). Analyzed with t-test.

(F) Representative flow plots of microglia and CD45$^{high}$ cells in the brains of the WT (*Csf1r*$^{+/+}$), *Csf1r*$^{\Delta FIRE/+}$, and *Csf1r*$^{\Delta FIRE/\Delta FIRE}$ mice. The number indicated the frequency of microglia of CD45$^{high}$ cells over all live, CD45$^+$, CD11b$^+$, Ly6G$^-$ cells.

(G) Percentage of GFP$^+$ cells in skull (blue), dura (orange) and blood (red) of *Csf1r*$^{\Delta FIRE/\Delta FIRE}$ mice joint with UBC-GFP mice (left), or received skull transplantation from UBC-GFP mice (right). The number indicated the number of cells recorded in the GFP$^+$ gate.

(H) Upper panel: representative flow gates of the percentage of $GFP^+$ cells of all $CD45^+$ cells in skull, dura and blood of a *Csf1r*$^{\Delta FIRE/\Delta FIRE}$ mouse paired from a UBC-GFP mouse. The number indicated the frequency of $GFP^+$ cells; Lower panel: representative flow gates of the percentage of $GFP^+$ cells of all $CD45^+$ cells in skull, dura and blood of a *Csf1r*$^{\Delta FIRE/\Delta FIRE}$ mouse after receiving skull cap from a UBC-GFP donor. The number indicated the frequency of $GFP^+$ cells.

(I) Whole-brain steromicroscopy imaging of $GFP^+$ cells on the brain surface of a *Csf1r*$^{\Delta FIRE/\Delta FIRE}$ mouse after skull transplantation from UBC-GFP donor mice.

(J) Widefield imaging of sagittal sections showing $GFP^+$ cells in brain parenchyma adjacent to transplant site.

(K) Experimental scheme of NaFl assay used to measure BBB permeability.

(L) Quantification of NaFl brain/blood ratio in age matched *Csf1r*$^{+/+}$ (N=9) and *Csf1r*$^{\Delta FIRE/\Delta FIRE}$ (N=7) mice. Data pooled from two independent experiments.

(M) Quantification of NaFl brain/blood ratio in age matched non-treated (NT, N=4) and PLX-3x treated (N=4) mice. Data collected from single experiment.
